# Modulation of noisy gene expression by general sequestration mechanisms

**DOI:** 10.64898/2026.07.30.741692

**Authors:** Ayan Biswas, Pavol Bokes, Abhyudai Singh

## Abstract

Sequestration of gene products through diverse mechanisms forms a fundamental layer of regulation in intracellular biochemical processes, including post-translational modification, promiscuous binding to genomic decoy sites, and partitioning into membraneless compartments formed through phase separation. Here, we develop a unified stochastic framework to quantify how such sequestration-type processes, when coupled to noisy gene expression, modulate cell-to-cell variation in protein levels. In this model, protein molecules reversibly switch between active (free) and inactive (sequestered) states, whose switching rates are arbitrary functions of the molecular counts. Using exact analytical calculations and the linear noise approximation, we derive expressions for the Fano factor of the active-protein level and identify fluctuation attenuation regimes in terms of the logarithmic sensitivities of the switching rates to protein abundances. We show that inactive-protein-dependent switching, of which genomic decoy binding is a natural example, can preserve Poisson-level fluctuations in the active-protein level under appropriate conditions. Enzymatic inactivation, a type of post-translational modification, emerges as a special case of active-protein-dependent sequestration, where greater responsiveness of the inactivation propensity attenuates active-protein fluctuations. In both decoy binding and enzymatic inactivation, protecting the inactive protein from decay lowers active-protein fluctuations. Finally, a noise-buffering regime associated with intracellular phase separation is recovered when the inactive-to-active switching rate depends inversely on the inactive-protein level. Together, the examples of genomic decoy binding, enzymatic inactivation, and intracellular phase separation suggest that noise buffering observed across diverse intracellular processes is rooted in a broader class of reversible sequestration mechanisms that attenuate protein-level fluctuations.

## I. INTRODUCTION

The functioning of a living cell is inseparable from stochasticity, which stems from the low abundance of biomolecules involved in key biochemical processes. The random nature of promoter switching via chromatin remodeling, as well as the probabilistic synthesis and degradation of mRNA and protein molecules, orchestrates marked intercellular expression variability despite identical genetic makeup at the cellular level [1–21]. Interestingly, organisms exploit this “isogenic heterogeneity” to hedge their bets for survival under adverse conditions. Such non-genetic phenotypic heterogeneity can compromise drug efficacy by enabling persistent or otherwise drug-tolerant states in bacterial and cancer-cell populations exposed to antibiotics and anticancer therapies, respectively [22–30]. Gene expression variability or “noise” facilitates cell-type diversification during early development [31–35], promotes “competence” for DNA uptake in *Bacillus subtilis* [36], enables the molecular switch for phage lysogeny and HIV-1 proviral latency [37–40], and also plays a pivotal role in mounting an effective immune response [41].

Previous work on *Saccharomyces cerevisiae* established that essential genes and genes coding for complex-forming proteins, e.g., proteasomal genes, show low expression noise. In contrast, genes that express proteins interacting with environmental signals, such as stress-related genes, are often highly noisy [7, 42, 43]. Given the importance of noise for diverse physiological functions and overall organismal fitness, cells have evolved elaborate regulatory mechanisms to control it. This is primarily because simple cellular signaling mechanisms cannot suppress noise to arbitrary precision due to demanding metabolic tradeoffs [44, 45]. In fact, a recent study on *Escherichia coli* suggests that noise regulation may be an important selectable organismal trait [46]. One of the prime examples of noise-attenuating mechanisms is negative autoregulation, where the transcription factor protein inhibits its own production via a nonlinear reaction rate, thereby providing stability against noise [47–53].

Cells also employ mechanisms other than transcriptional regulation to control biochemical noise. Protein sequestration, which generates an ultrasensitive “all-or-none” response in various cell fate decisions, plays a significant role in modulating cellular stochasticity [54–56]. Ultrasensitivity can create a “low-pass filtering” mechanism to limit the propagation of input signal fluctuations along transcriptional cascades [57]. Depending upon the kinetic timescales and cooperativity strength, sequestration mechanisms can buffer against noise and change the nature of protein distribution [58]. ChIP-chip and ChIP-seq analyses have identified genome-wide binding sites for transcription factors that vastly outnumber known binding sites linked to direct regulatory processes [59]. The muscle differentiation factor protein MyoD constitutively binds to thousands of sites, resulting in epigenetic modifications in addition to direct muscle gene regulation [60]. By contrast, the maternal and gap transcription factors responsible for initial anterior-posterior patterning in *Drosophila melanogaster* development have been found to bind to many genes at low levels with no functional outcomes [61]. Theoretical studies have considered the possibility that genomic nonregulatory “decoy” sites may protect transcription factors from degradation by sequestering the proteins. This process suppresses the noise via attenuating the correlations in the protein pool [62]. In fact, for an autoregulated gene, these decoy sites can decrease the noise in the free (unbound) protein pool to the Poisson level [63]. Another interesting class of mechanisms involves post-translational modifications, e.g., enzymatic reactions, as observed in the signal-amplifying MAP kinase cascades [64]. Other examples include phosphorylation-driven inactivation of anti-anti-sigma factor during sporulation in *Bacillus subtilis* [65] and inactivation of repressors by antirepressor binding in *Salmonella* phage P22 [66]. However, the role of noise in the latter cases is not well characterized. Recent studies have also pointed out that phase separation of biomolecules in membraneless compartments buffers against noise [67, 68].

In this paper, we propose a unified “sequestration-type” framework that can characterize the noisy behavior of proteins switching reversibly between active and inactive states. A key distinction between the two states in our model is that an inactive protein may be protected from decay, whereas the active protein is not. The active-to-inactive and inactive-to-active state switching is analogous to the sequestration and dissociation reactions, respectively. The switching rates are taken to be arbitrary functions of the active- and inactive-protein levels, subject to the existence of a unique stable steady state. Interestingly, when protein synthesis is non-bursty, the switching rates depend only on the inactive-protein level, and the inactive protein is protected from decay, we find that the active-protein distribution remains Poisson and is unaffected by the sequestration mechanism. Furthermore, we quantify the steady-state Fano factor of the active-protein level as a function of the switching-rate sensitivities and demarcate the regimes in which active-protein fluctuations are attenuated. We use the level of active-protein fluctuations from the production-and-degradation process (i.e., without sequestration) as our primary reference for identifying effective suppression of fluctuations. Depending on whether the inactivation protects the protein from decay, the quantitative nature of fluctuation attenuation differs.

We connect the analytical findings emanating from our generalized stochastic framework to examples of post-translational modification of proteins and intracellular phase separation with the support of stochastic simulations. The generalized models corresponding to these special cases involve an active-protein-dependent sequestration rate (enzymatic inactivation of proteins) and an inactive-protein-dependent dissociation rate (intracellular phase separation). We also map the fluctuation-attenuating effects arising from the arbitrarily chosen inactive-protein-dependent sequestration rate to the specific case of sequestration of transcription factor protein at genomic decoy sites. In all of these cases, we focus on identifying the fluctuation characteristics of the active-protein level from arbitrary sequestration or dissociation mechanisms. The low fluctuation level in the free (active) protein pool when the inactive protein is protected from decay in the decoy binding case also appears in the case of enzymatic inactivation of proteins, suggesting a shared underlying mechanism. This possibility also exists in the case of phase separation, as new theoretical results confirm that noise attenuation can be achieved independent of concentration buffering in biomolecular condensates [69]. Our findings for the generalized inactive-protein-driven dissociation reaction support the view that fluctuation attenuation can be observed for a broad class of intracellular processes and is not unique to phase separation.

The paper is structured as follows. In Sec. II, we formulate our stochastic model and discuss different sequestration-type mechanisms. We provide some exact results in the special case of non-bursty protein synthesis in Sec. III A. The corresponding extension to protein multimerization is provided in Appendix A. Sec. III B deals with the case of sequestration with constant switching rates. Details of the associated analytical calculations are provided in Appendix B. We characterize the profiles of active-protein fluctuations when the active and inactive proteins control the state switching in Sec. III C and Sec. III D, respectively. The analyses regarding the existence of unique stable steady states in these cases are presented in Appendix C. Finally, in Sec. IV, we summarize our findings and point out potential extensions of the current theoretical framework.

## II. MODEL FORMULATION AND EXAMPLE CASES OF INTRACELLULAR SEQUESTRATION-TYPE MECHANISMS

In our stochastic gene expression model, protein molecules (*X*) are synthesized from the active genetic promoter in bursts that arrive as a Poisson process with rate *k*_*x*_. The burst sizes *B*_*x*_ are independent and identically distributed, non-negative integer-valued random variables following an arbitrary probability mass function [8, 10, 53, 70, 71]

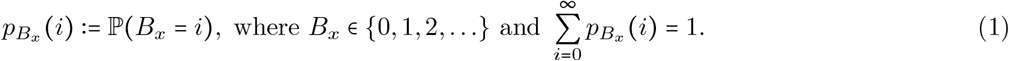

In the stochastic description, each biochemical reaction is characterized by a propensity function, defined such that the probability of its occurrence in the next infinitesimal time interval (*t, t* + *dt*] is given by the propensity evaluated at the current molecular state, multiplied by *dt* [72]. Hence, in an infinitesimal time interval (*t, t* + *dt*], the propensity of protein synthesis of a burst size *i* is 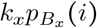 and the corresponding transition probability is 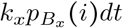. These protein molecules are also subject to decay with rate *γ*_*x*_ and propensity *γ*_*x*_*x*(*t*), where *x*(*t*) is the number of protein molecules at time *t*. Thus, 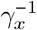 is the characteristic lifetime associated with decay of the active protein. Furthermore, an active protein molecule may be sequestered and thereby enter an inactive state, denoted by *Y*, whose copy number at time *t* is *y t* . The sequestration rate generally depends on the protein levels in both states and is denoted by *k*_s_ (*x, y*). Here, one active protein molecule is sequestered in each reaction, so the corresponding propensity is *k*_*s*_ (*x, y*) *x*. More generally, multiple active protein molecules may be sequestered cooperatively to form an inactive complex. One prominent example is when bacteria downregulate ribosomal activity by dimerizing 70S ribosomes into inactive complexes [73, 74]. The sequestered (inactive) proteins can also switch back to their active state with a protein-level-dependent dissociation rate *k*_*d*_ (*x, y*) and propensity *k*_*d*_ (*x, y*) *y*. Lastly, the inactive proteins decay with a propensity *γ*_*y*_*y*, where *γ*_*y*_ is the decay rate, which is, in general, different from *γ*_*x*_. The relative decay rate of the inactive protein is defined as the dimensionless ratio *β γ*_*y*_ */ γ*_*x*_. The limiting case *β =* 0 represents a protected inactive state, where the sequestered proteins do not decay, whereas *β* = 1 corresponds to inactive proteins decaying at the same rate as active proteins. Thus, *β* determines whether sequestration creates a stabilized storage reservoir or merely an alternative molecular state without additional protection from decay. Such distinctions between free and bound molecular states arise naturally in transcription-factor decoy binding, where bound proteins may be protected from decay, although context-dependent decay of the bound protein has also been considered [62, 75, 76]. In each of these biochemical reactions, molecular counts of the participating species change by integer amounts with the probability defined by the respective propensity functions [72, 77]. The reaction schemes are described in Table I and illustrated in Fig. 1.

**FIG. 1.**
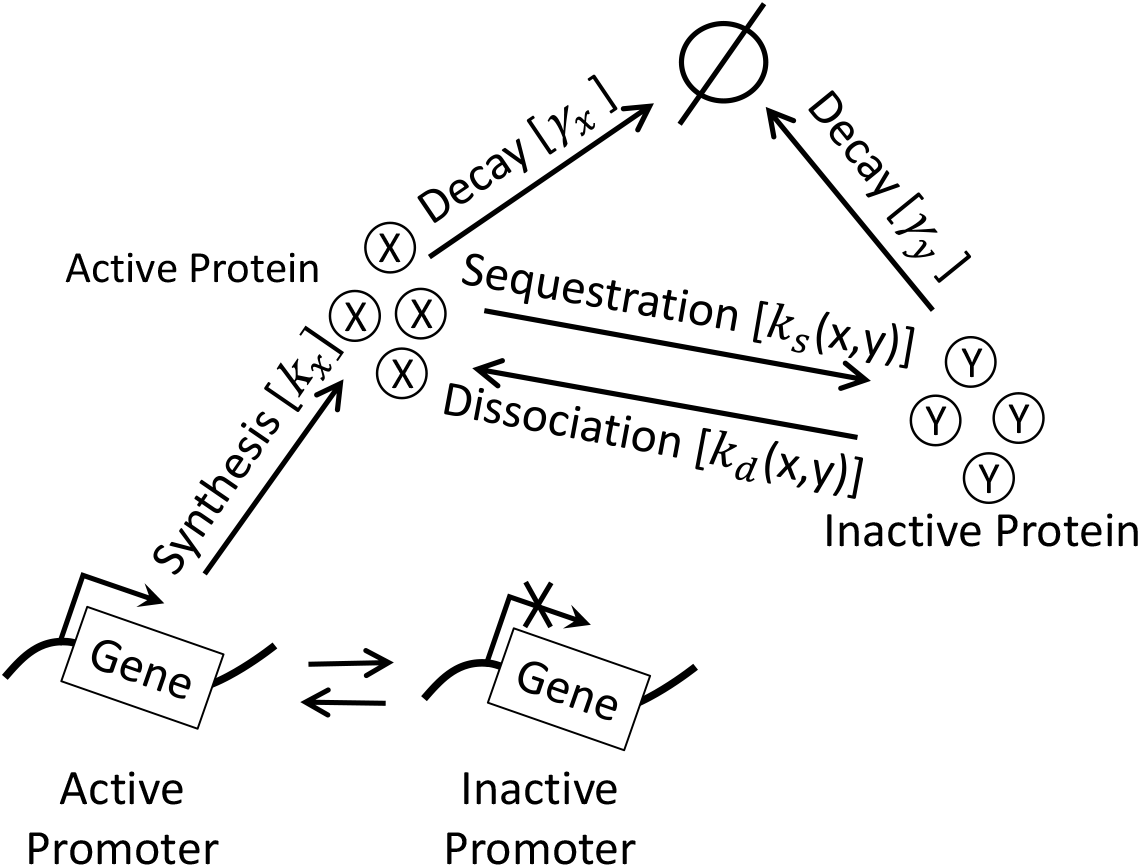
Schematic representation of bursty gene expression coupled to molecular sequestration and dissociation. Bursty protein synthesis is modeled as an effective coarse-grained process that captures the influence of underlying promoter-state dynamics without explicitly modeling promoter switching. The synthesized proteins reversibly switch between active and inactive states through sequestration and dissociation, and proteins in both states may undergo decay. The rates of the individual biochemical reactions are indicated in brackets. The sequestration and dissociation rates are arbitrary functions of the active- and inactive-protein levels.

**TABLE I.** Model for bursty gene expression with sequestration and dissociation kinetics. Proteins are synthesized from an active promoter in bursts at a rate *k*, with a burst of size *i* occurring with probability 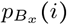. Active proteins are sequestered (inactivated) and can revert to their active state (dissociation/reactivation). *x* (*t*) and *y* (*t*) are the active and inactive protein copy numbers, respectively. The sequestration and dissociation rates are arbitrary functions of protein levels, denoted by *k*_*s*_ (*x, y*) and *k*_*d*_ (*x, y*), respectively. Active and inactive proteins decay at rates *γ*_*x*_ and *γ*_*y*_, respectively.

| Biochemical reactions | Changes in species counts | Probability of reaction in the time interval $(t, t + dt]$ |
| --- | --- | --- |
| Protein synthesis | $x \mapsto x + i$<br>$i \in \{0, 1, 2, \dots\}$ | $k_x p_{B_x}(i) dt$<br>$p_{B_x}(i) := \mathbb{P}(B_x = i)$ |
| Sequestration (active-to-inactive state switching) | $x \mapsto x - 1$<br>$y \mapsto y + 1$ | $k_s(x, y) x dt$ |
| Dissociation (inactive-to-active state switching) | $x \mapsto x + 1$<br>$y \mapsto y - 1$ | $k_d(x, y) y dt$ |
| Decay of active protein | $x \mapsto x - 1$ | $\gamma_x x dt$ |
| Decay of inactive protein | $y \mapsto y - 1$ | $\gamma_y y dt$ |

Our generalized stochastic framework can be used to quantify fluctuations in protein levels across various biochemical processes that involve switching between two alternative activity states of proteins. The active- and inactive-protein copy numbers are denoted by *x* and *y*, respectively; we refer to these molecular abundances more generally as protein levels.

- One such biochemical process that can be readily modeled using our framework is the post-translational modification of proteins. In our model, these reactions can be captured with the Michaelis-Menten-type sequestration rate,

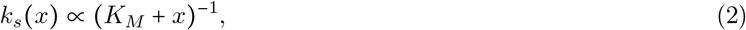

where *K*_*M*_ is the Michaelis constant. The reactivation process may be characterized by a protein-independent (constant) dissociation rate.
- Genomic decoy sites can protect transcription factor proteins from decay by sequestering them. These noncoding sites also buffer protein-level fluctuations [62, 63, 78], and decoy–transcription-factor complexes can improve the precision of biochemical-event timing [79, 80]. Here, *y* denotes the number of bound proteins, or equivalently the number of occupied decoy sites, so that *y* ∈ {0, 1, …, *M*} . The sequestration rate depends on the number of unoccupied sites according to

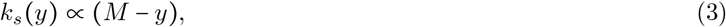

where *M* is the total number of genomic decoy sites [76]. The dissociation rate may again be taken to be independent of protein levels.
- As a final example, we consider a state-switching scenario akin to intracellular phase separation in eukaryotes, in which proteins and nucleic acids form membraneless compartments [81]. The sequestration and dissociation rates depend on the levels of biochemical species in the dilute phase (*x*) and droplet phase (*y*) as [67]

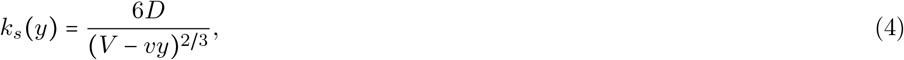

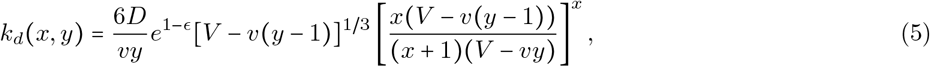

respectively. Here, *D* is the diffusion constant, *V* is the volume of the system, and *v* is the volume of a protein molecule. Hence, *vy* denotes the volume of the membraneless phase-separated compartment (droplet phase). *ϵ* is a dimensionless interaction parameter. For small nonzero *y* with *vy* / *V* ≪ 1, a dilute active-protein volume fraction satisfying *xv*/*V* ≪ 1, and a sufficiently large active-protein count *x* such that (*x*/(*x* + 1))^*x*^ ≈ *e*^−1^, Eqs. (4)–(5) reduce to

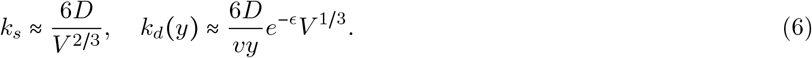

Hence, under these limiting conditions, the sequestration propensity is first order in the active-protein level. For *y* > 0, the dissociation propensity *k*_*d*_ (*y*) *y* is effectively zeroth order, whereas it is defined to be zero at *y =* 0 because no inactive protein is available for dissociation. The simplified rates therefore correspond to an effectively constant sequestration rate and a dissociation rate that decreases inversely with the inactive-protein level. Table II summarizes the switching rate forms for these biochemical processes.

**TABLE II.**
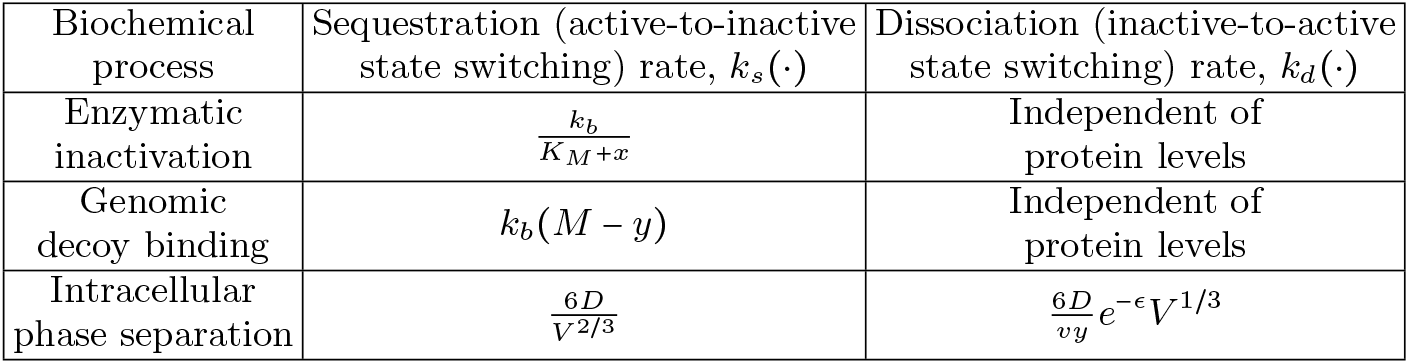
Functional forms of the switching rates in different biochemical processes. For enzymatic inactivation, *k*_*b*_ and *K*_*M*_ denote the maximal inactivation capacity of the enzyme system and the Michaelis constant, respectively. For genomic decoy binding, *k*_*b*_ and *M* denote the binding-rate parameter and the total number of genomic decoy sites, respectively. In the context of phase separation, *D* is the diffusion constant, *V* is the system volume, *v* is the volume of the protein molecule, and *ϵ* is a dimensionless interaction parameter. For the phase-separation rates, we consider the limiting regime *vy / V* ≪ 1, *xv / V* ≪ 1, and *x* ≫ 1 in Eqs. (4)–(5).

## III. RESULTS AND DISCUSSION

### A. Sequestration with non-bursty protein synthesis and protected inactive protein

Here, we consider a special case of the reaction kinetics in Table I: protein synthesis is non-bursty, i.e., *B*_*x*_ = 1 with probability 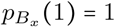. The switching rates are arbitrary functions of the inactive-protein level, i.e., *k*_*s*_ (*x, y*) = *k*_*s*_ (*y*) and *k*_*d*_(*x, y*) = *k*_*d*_(*y*). The inactive protein is protected from decay (*γ*_*y*_ = 0). The steady-state solution to the corresponding master equation is obtained in a separable form [82–85]:

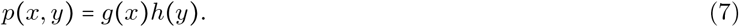

The marginal steady-state distribution of the active protein is Poisson [83],

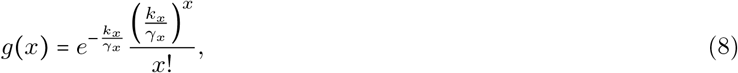

with the mean

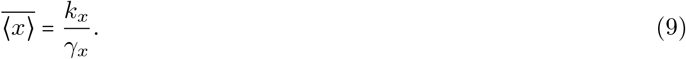

Hence, the inactive-protein-mediated switching does not alter the Poisson distribution of the active protein under non-bursty synthesis and first-order decay. The inactive-protein distribution is non-Poisson in general,

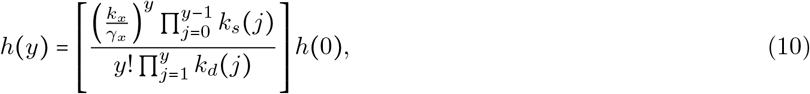

where *h*(0) is determined from the normalization condition:

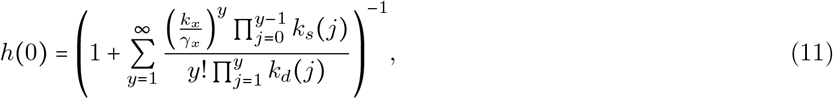

provided that the normalization sum converges. The structure of Eqs. (10)–(11) can be interpreted in terms of an effective birth–death process for the inactive-protein count: transitions *y* ↦ *y +* 1 occur with propensity (*k*_*x*_ */ γ*_*x*_ ) *k*_*s*_ (*y*), whereas transitions *y* ↦ *y* − 1 occur with propensity *yk*_*d*_ (*y*). Biochemically, these transitions represent sequestration and dissociation, respectively. When *k*_*s*_(*y*) = *k*_*s*_ and *k*_*d*_(*y*) = *k*_*d*_ are independent of protein levels, the resulting inactive-protein distribution is Poisson, with mean

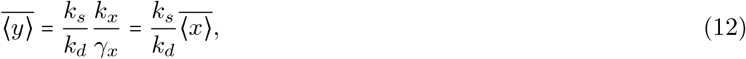

where in the last equality, we have used Eq. (9).

Considering *M* genomic decoy sites to which free proteins bind, with *y* denoting the number of bound proteins, or equivalently the number of occupied decoy sites, the sequestration rate depends on the number of free decoy sites as *k*_*s*_ (*y*) ∝ (*M* − *y*), while the dissociation rate *k*_*d*_ is constant. For this choice of switching rates, Eqs. (10)–(11) recover the previously reported stationary result for reversible transcription-factor binding to a finite number of downstream DNA sites [83]: the free transcription-factor count is Poisson distributed, independently of the downstream process, whereas the number of transcription-factor–DNA complexes follows a binomial distribution.

Our framework can be generalized to cooperative sequestration, in which multiple active protein molecules are sequestered together to form an inactive complex. The corresponding results are provided in Appendix A. We recover the Poisson distribution of the active-protein count in Eqs. (A12)–(A13). The inactive-complex distribution retains the same product-form structure as the inactive-protein distribution in the monomeric case, as shown in Eqs. (A15)–(A16). When the switching rates are constant, the inactive-complex count follows a Poisson distribution, similar to the inactive-protein count in the absence of multimerization. Eq. (A17) gives the corresponding mean inactive-complex count, generalizing the mean inactive-protein level in Eq. (12). These multimerization-specific results may be relevant to bacteria that downregulate ribosomal activity by dimerizing 70S ribosomes into inactive complexes [73, 74].

### B. Sequestration with constant switching rates

After considering the non-bursty, protected-inactive-state limit in Sec. III A, we now return to the general model with bursty protein synthesis introduced in Sec. II and summarized in Table I. We consider a simple switching mechanism in which the sequestration and dissociation rates are independent of protein levels, i.e., *k*_*s*_ (*x, y*) *= k*_*s*_ and *k*_*d*_ (*x, y*) = *k*_*d*_ . The active protein undergoes decay, while the inactive protein may either be protected or decay, as determined by the relative decay rate *β* = *γ*_*y*_/*γ*_*x*_. We use moment dynamics [86–90] to obtain the first two steady-state moments of the active-protein copy number and compute its Fano factor,

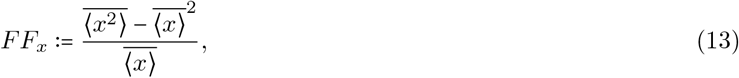

which measures active-protein fluctuations as the variance relative to the mean. The corresponding analytical derivation is provided in Appendix B. The unique stable steady-state mean protein levels are given by

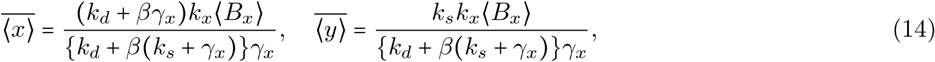

for burst sizes (*B*_*x*_) distributed following an arbitrary probability mass function 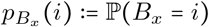, *i* ∈ {0, 1, 2, … } with its first-order moment defined as

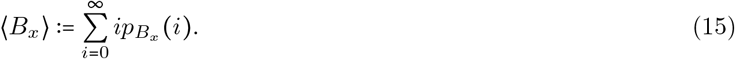

The mean active-protein level in the absence of sequestration is obtained by substituting *k*_*s*_ *=* 0 in the first expression in Eq. (14):

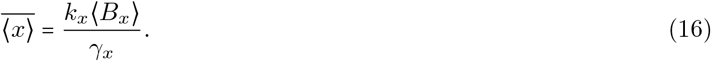

As mentioned in Sec. II, we focus on two limiting cases of inactive protein decay: *β* = 0 (inactive protein is protected from decay) and *β* = 1 (inactive protein decays at the same rate as the active protein).

We use the expression of 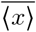 from Eq. (14) and that of 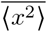 from Eq. (B5) in Eq. (13) to write the Fano factor in an exact form:

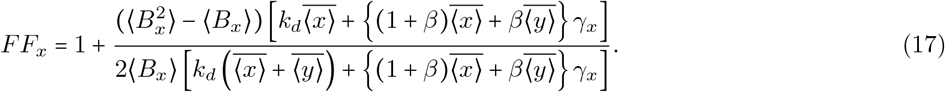

Here, 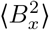 represents the second-order moment of the burst size distribution,

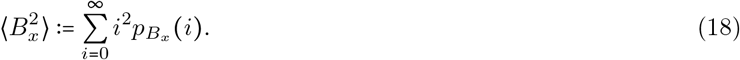

In the absence of sequestration 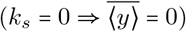, Eq. (17) reduces to [76],

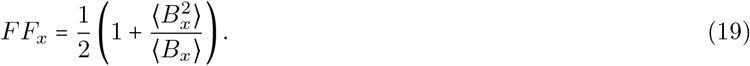

For the non-bursty protein synthesis, i.e., burst size *B*_*x*_ = 1 with probability 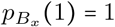, we have 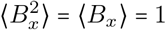. In this situation, Eq. (19) reproduces the standard Poisson result,

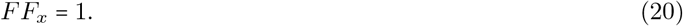

In fact, non-bursty protein synthesis, even in the presence of sequestration, keeps the Fano factor at the Poisson level for both *β =* 0 and 1, as can be verified from Eq. (17).

When the sequestration and dissociation rates are much faster than the protein decay rate and the steady-state mean protein levels are held fixed, Eq. (17) takes a simplified form. This corresponds to taking *k*_d_ → ∞, with *k*_*s*_ → ∞ accordingly in Eq. (17), while keeping 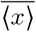 and 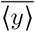 fixed. In this fast-switching limit, the Fano factor becomes

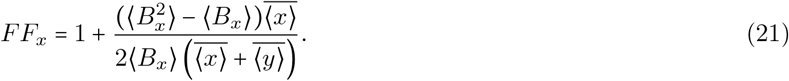

Experimental evidence suggests that protein burst sizes are often independent and identically distributed, non-negative integer-valued random variables (*B*_*x*_) that follow a geometric probability mass function [8, 91–93]:

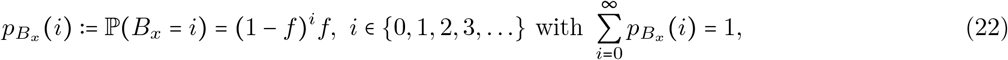

where

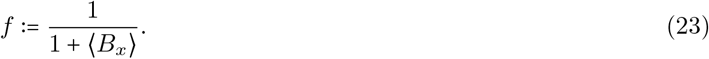

The first- and second-order moments of the geometric burst size distribution are related via

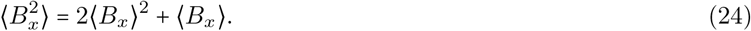

We use Eq. (24) in our general expression from Eq. (17) to obtain the Fano factor in terms of the average burst size ⟨*B*_*x*_⟩ :

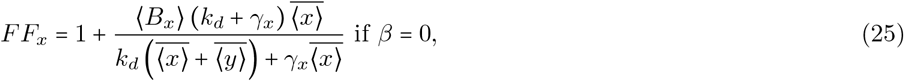

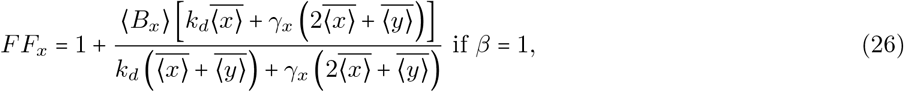

which indicates attenuation in active-protein fluctuations with increasing dissociation rate *k*_*d*_, when the steady-state mean protein levels are held fixed. Similarly, for geometrically distributed burst sizes, the expression of *FF*_*x*_ in the fast-switching limit in Eq. (21) takes the for

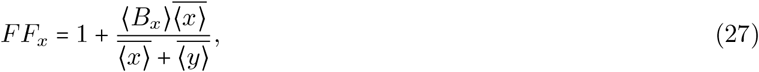

which, unlike Eqs. (25) and (26), is independent of *β*. Lastly, the case without sequestration in Eq. (19) produces

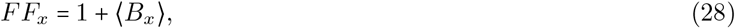

for geometrically distributed burst sizes.

To probe the effect of sequestration on active-protein fluctuations, we use the ratio of Fano factors with and without sequestration. For this, *FF*_*x*_ from Eqs. (25) and (26) (with sequestration) is normalized by the *FF*_*x*_ from Eq. (28) (without sequestration) and is plotted in Fig. 2 as a function of the dissociation rate *k*_*d*_. *FF*_*x*_ for the fast-switching case in Eq. (27) is similarly normalized and plotted for comparison. Looking at the Fano factor ratios in Fig. 2, we note that the fluctuations in the active-protein level due to sequestration with and without the fast-switching condition do not exceed the corresponding fluctuations without sequestration and are strictly lower for *k*_*d*_ > 0. The active-protein fluctuations are higher for *β* = 1 than for *β* = 0. The fluctuations in the fast-switching limit are the least of all three cases.

**FIG. 2.**
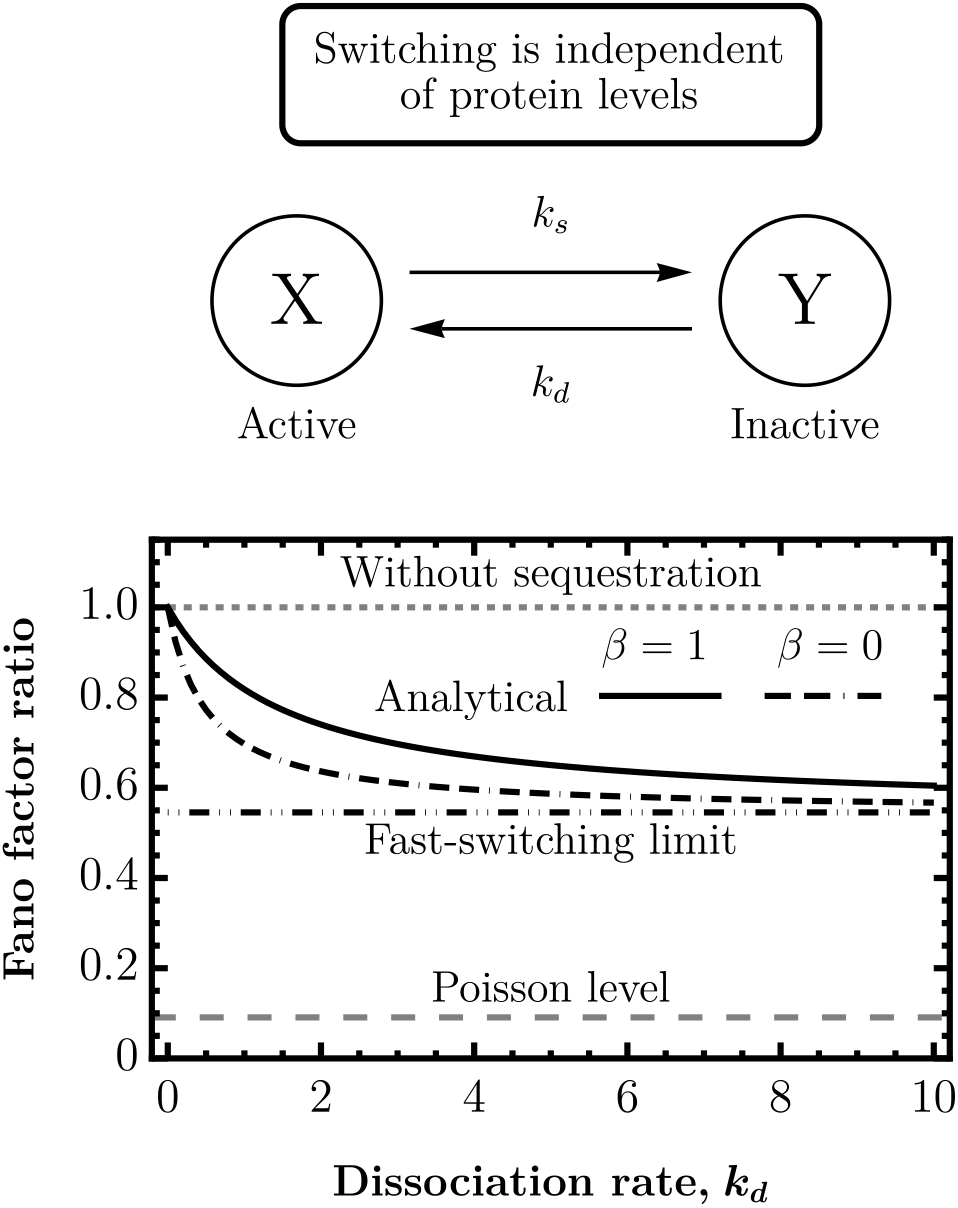
Active-protein fluctuations are attenuated with increasing dissociation rate when switching rates are independent of protein levels. The ratio of the active-protein Fano factor under sequestration, *FF*_*x*_ from Eqs. (25) and (26), to the Fano factor without sequestration, *FF*_*x*_ from Eq. (28), is plotted as a function of the dissociation rate *k*_*d*_. The corresponding Fano factor ratio in the fast-switching limit, calculated using Eqs. (27) and (28), is also shown. The “Without sequestration” and “Poisson level” reference values, the latter given by the ratio of *FF*_*x*_ in Eq. (20) to *FF*_*x*_ in Eq. (28), are indicated by short- and long-dashed gray lines, respectively. Parameters: 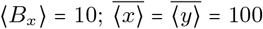 copies; *γ*_*x*_ = 1 and *γ*_*y*_ = 0 for *β* = 0; and *γ*_*x*_ = *γ*_*y*_ = 1 for *β* = 1. The protein synthesis rate *k*_*x*_ is chosen to achieve the prescribed steady-state protein levels, while the sequestration rate *k*_*s*_ is adjusted as *k*_*d*_ is varied to keep these levels fixed.

### C. Active protein controls sequestration rate

In situations such as enzymatically driven protein inactivation, the active-to-inactive switching rate depends only on the active-protein level, *k*_*s*_ (*x*), whereas the dissociation rate *k*_*d*_ is constant. To model such sequestration mechanisms, we take the sequestration rate to have the form

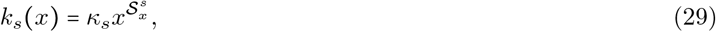

where *κ*_*s*_ is the rate parameter and 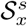 denotes the logarithmic sensitivity of the sequestration rate to the active-protein concentration. We apply the linear noise approximation (LNA) to the nonlinear sequestration propensity 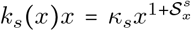 to obtain the active-protein Fano factor in closed form. In applying the LNA, we assume that fluctuations in the protein levels are small relative to their respective deterministic steady states [94–97]. The general deterministic reaction-rate equations, corresponding to the stochastic reaction scheme in Table I, are given by Eqs. (C1) and (C2). The steady-state protein concentrations are denoted by 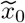 and 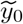. Conditions ensuring monostability of the deterministic system under active-protein-dependent sequestration are discussed in Appendix C 1. The sensitivity evaluated at steady state is defined as:

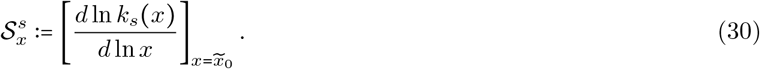

Here and below, the power-law switching rates are used as local parametrizations about positive steady states; for exact stochastic dynamics, their zero-copy boundaries are specified by the corresponding biochemical propensities, with any reaction consuming an absent species assigned zero propensity.

For *β =* 0, a unique steady state exists for any 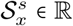 provided *k*_*d*_ > 0. The corresponding protein concentrations are

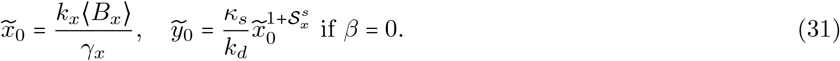

The steady-state active-protein concentration is the same as in the case without sequestration (see Eq. (16)). With 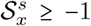 the unique steady state is guaranteed to be locally asymptotically stable. For *β =* 1, the steady-state active-protein concentration does not, in general, admit a closed-form expression but satisfies

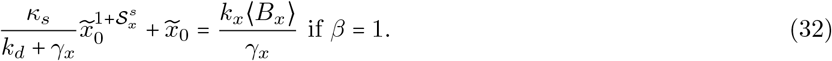

For 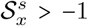, or equivalently 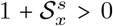, the left-hand side of Eq. (32) is continuous on [0, ∞), strictly increasing on (0, ∞), approaches zero as 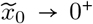,, and diverges as 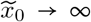. Hence, for *k*_*x*_⟨*B*_*x*_⟩/*γ*_*x*_ > 0, Eq. (32) admits a unique positive root, without any further constraints on the other parameters. The corresponding unique steady-state inactive-protein concentration is

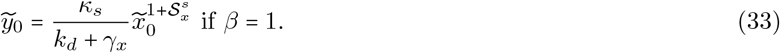

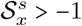 ensures the local asymptotic stability of steady state in the *β =* 1 case. The stability analysis is presented in Appendix C 1.

We next use the LNA and expand the nonlinear sequestration propensity around the unique and stable steady state 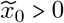 and retain terms up to the linear order:

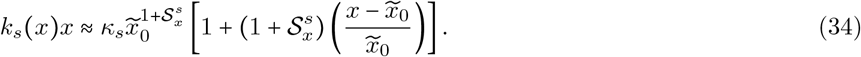

We incorporate this linearized sequestration propensity in the moment dynamics equation [88–90] and evaluate the required steady-state moments. The steady-state mean protein levels are

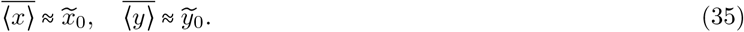

For geometrically distributed burst sizes 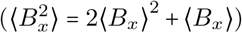, the following closed-form Fano factor expressions for the active protein are obtained in the fast-switching limit.

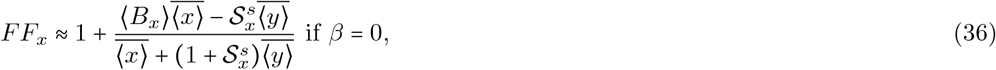

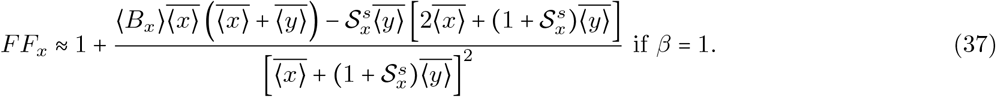

Both Eqs. (36) and (37) indicate attenuation in the active-protein fluctuations with an increasing sequestration sensitivity to the active-protein level 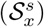.

The Fano factors from Eqs. (36), (37) are normalized by the Fano factor without sequestration (*FF*_*x*_ = 1 + ⟨*B*_*x*_⟩ from Eq. (28)). The Fano factor ratios are plotted in Fig. 3A. We observe that an increasing sensitivity of the sequestration rate to the active-protein level leads to attenuation of fluctuations in the active-protein level below the level of fluctuations without sequestration. For the non-bursty gene expression (*B*_*x*_ = 1 with probability 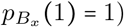, we have 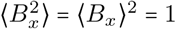. Furthermore, if inactive proteins are protected from decay (*β* = 0), the Fano factor has the form

**FIG. 3.**
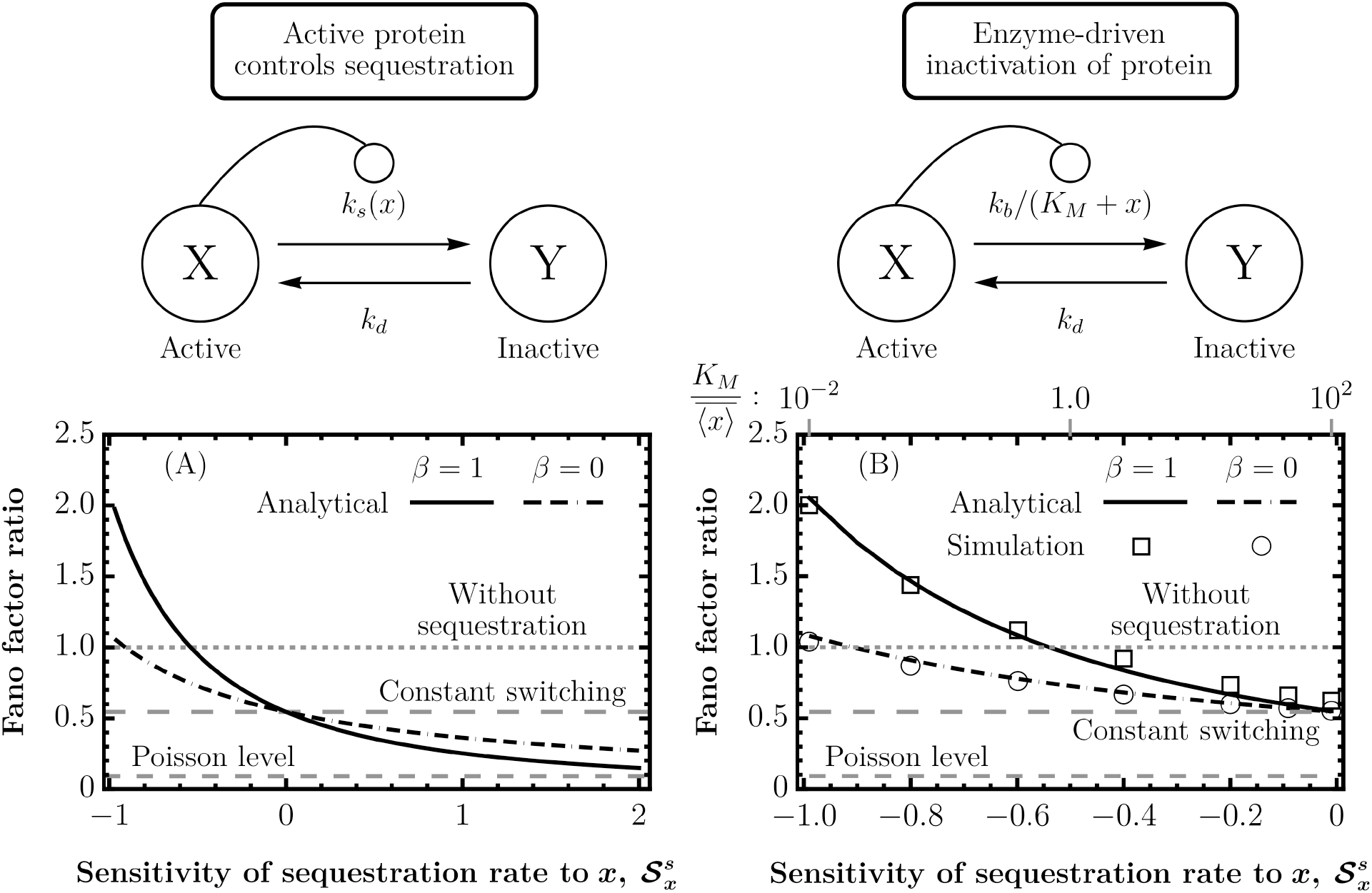
Active-protein fluctuations are attenuated with increasing sensitivity of the sequestration rate to the active-protein level. (A) The ratios of the active-protein Fano factors under sequestration, *FF*_*x*_ from Eqs. (36) and (37), to the Fano factor without sequestration, *FF*_*x*_ from Eq. (28), are plotted as functions of the sensitivity ^*s*^. Gray dashed lines indicate the “Without sequestration”, “Constant switching” (the ratio of *FF*_*x*_ in Eq. (27) to *FF*_*x*_ in Eq. (28)), and “Poisson level” (the ratio of *FF*_*x*_ in Eq. (20) to *FF*_*x*_ in Eq. (28)) reference values. The “Constant switching” reference corresponds to the”Fast-switching limit” line in Fig. 2. (B) An example of post-translational modification in which the protein is enzymatically inactivated. The sequestration rate is *k*_*s*_ (*x*) *= k*_*b*_ */* (*K*_*M*_ *+ x*), where *k*_*b*_ is the maximal inactivation capacity of the enzyme system, and the dissociation rate *k*_*d*_ is constant. The Michaelis constant *K*_*M*_ is varied from 1 to 10^4^ (gray ticks along the top axis), and the sensitivity ^*s*^ is calculated using Eq. (43). Analytical lines show the corresponding Fano factor ratios in the fast-switching limit; symbols show stochastic-simulation estimates [98, 99] from 10^4^ independent endpoint samples at steady state. Parameters: 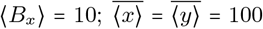 copies; *γ*_*x*_ = 1 and *γ*_*y*_ = 0 for *β* = 0; and *γ*_*x*_ = *γ*_*y*_ = 1 for *β* = 1. The stochastic simulations use the finite switching rate *k*_*d*_ = 10*γ*_*x*_. Other parameters are adjusted to keep the steady-state protein levels fixed as 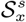 is varied.

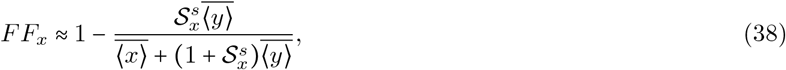

in the fast-switching limit. Thereby, we recover the monotonically decreasing trend of the active-protein fluctuations with an increasing sensitivity 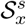 as reported in [100]. From Eq. (38), we notice that fluctuations in the active-protein level are super-Poisson for negative sensitivity and go to the sub-Poisson regime as the sensitivity becomes positive. In contrast, over the sensitivity range shown in Fig. 3A for bursty protein synthesis, the active-protein fluctuations remain in the super-Poisson regime.

The model can be adapted to analyze fluctuations in active-protein levels regulated by post-translational modifications, such as enzymatically driven inactivation. To this end, we use the sequestration rate form

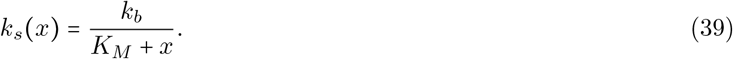

Thus, the sequestration propensity becomes

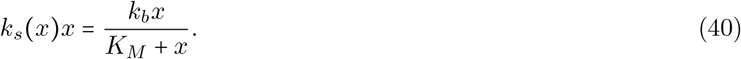

Here, the Michaelis constant *K*_*M*_ denotes the active-protein level at which the sequestration propensity reaches half of its maximal value, *k*_*b*_ */* 2. It is defined as

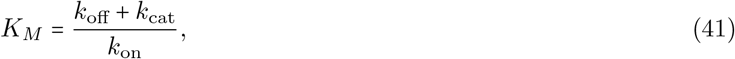

where *k*_on_ and *k*_off_ are the binding and unbinding rate constants for reactions involving the enzyme and the active protein. *k*_cat_ is the turnover number of the enzyme and signifies how fast the enzyme can convert the active protein into its inactive form when the active protein is abundant. The maximal inactivation capacity of the enzyme system is denoted by

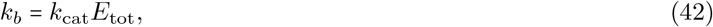

where *E*_tot_ is the total enzyme concentration, including both free and complex-bound forms. The dissociation rate *k*_*d*_ for the inactive-to-active state switching is independent of protein levels.

Under the standard quasi-steady-state assumption, the enzyme-active-protein complex rapidly relaxes to its quasi-steady state at each moment relative to the active-protein dynamics, yielding the Michaelis-Menten form in Eq. (40). In our parameter sweep, *K*_*M*_ and *k*_*b*_ are increased simultaneously so that the steady-state active- and inactive-protein levels remain fixed. The Michaelis-Menten form of the sequestration propensity is most directly justified when the enzyme level is sufficiently low that the enzyme-active-protein complex level is negligible relative to the active-protein level [101]. When enzyme and protein levels are comparable, as in protein interaction networks, a modified propensity based on an appropriate revised quasi-steady-state approximation should instead be used [102]. Even when the deterministic quasi-steady-state approximation is valid, however, the corresponding stochastic reduction can yield inaccurate estimates of intrinsic fluctuations, particularly at low molecular copy numbers [103]. From the definition in Eq. (30) with the substitution 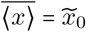, the sensitivity of the sequestration rate turns out to be

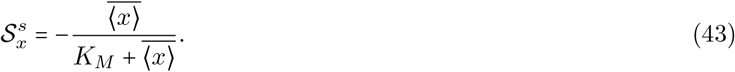

Hence, for fixed 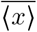, changing *K*_*M*_ alters the sensitivity 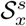, and therefore modifies *FF*_*x*_. This enzymatic inactivation example was previously investigated for the non-bursty case with *β* = 0, without imposing the fast-switching condition. In that case, we found that 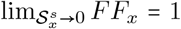 [100], consistent with Eq. (38). For the present case, the Fano factor ratios in the fast-switching limit are computed using Eqs. (36), (37), and (28), and are shown as lines in Fig. 3B. The analytical results are supported by stochastic simulation algorithm results [98, 99], shown as symbols.

An increasing Michaelis constant or half-saturation level (*K*_M_ ) increases the sensitivity 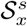 (makes it less negative) via Eq. (43), thereby attenuating active-protein fluctuations. When 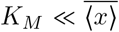, the steady-state sequestration (inactivation) propensity satisfies 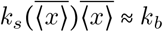, reflecting the nearly protein-independent behavior of the inactivation propensity in the saturated regime. At the opposite extreme, when 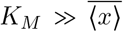, we get 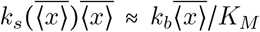, reflecting an approximately linear dependence of the inactivation propensity on the active-protein level.

Thus, with increasing *K*_*M*_, the enzyme system shifts from a saturated, approximately zero-order regime to a less saturated, approximately first-order regime. Because *k*_*b*_ is simultaneously increased to maintain the same steady-state protein levels, the maximal inactivation capacity of the enzyme also increases. The responsiveness of the complete sequestration (inactivation) propensity is best captured by its logarithmic sensitivity, 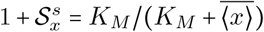, which increases monotonically with *K*_*M*_ . Consequently, the inactivation propensity responds more strongly whenever the active-protein level deviates from its steady state, producing an implicit restoring mechanism analogous to negative feedback. This increased responsiveness attenuates fluctuations in the active-protein level.

In Fig. 3B, when both protein species decay at the same rate (*β =* 1), the fluctuations exceed the “Without sequestration” level over a substantial range of sensitivity. By contrast, when the inactive protein is protected from decay (*β* = 0), the fluctuations remain predominantly below this level. As the Michaelis constant increases and the sensitivity approaches zero, the active-protein fluctuations approach the “Constant switching” limit for both *β* = 0 and *β* = 1. For nonzero sensitivity, our results show that protecting the inactive protein from decay (*β* = 0) reduces fluctuations in the active-protein level. The active protein trajectories and probability density histograms in Fig. 4 further demonstrate that increasing the half-saturation level, and hence increasing sensitivity, attenuates fluctuations.

**FIG. 4.**
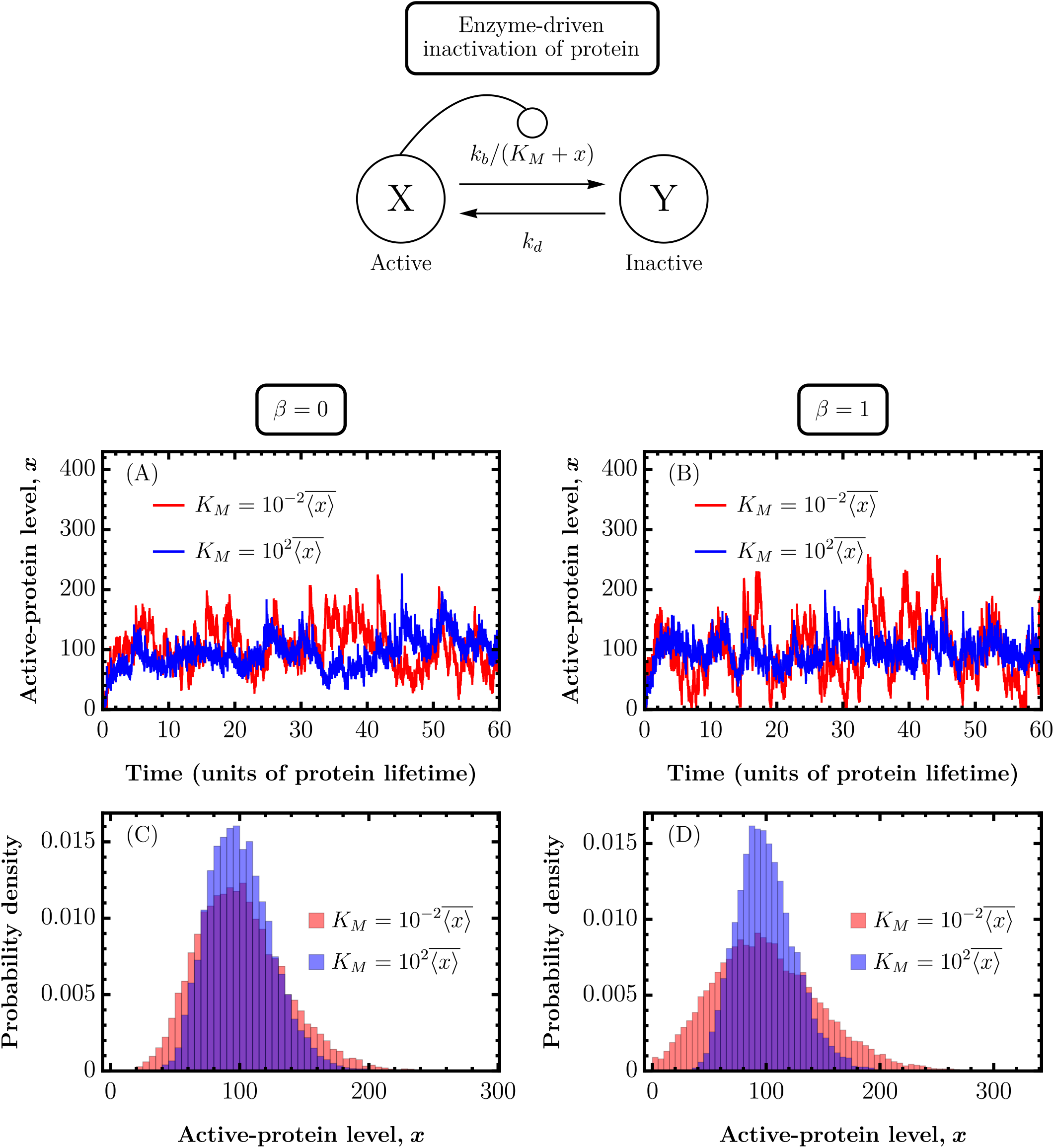
Active-protein fluctuations are attenuated by enzymatic inactivation (continued from Fig. 3B). Here, *k*_*s*_ (*x*) *= k*_*b*_ */* (*K*_*M*_ *+ x*), where *k*_*b*_ is the maximal inactivation capacity of the enzyme system and *K*_*M*_ is the Michaelis constant; *k*_*d*_ is constant. At fixed steady-state protein levels, *k*_*b*_ increases with *K*_*M*_ . (A, C) Sample trajectories and probability density histograms from stochastic simulations [98, 99], respectively, for different values of *K*_*M*_ when *β* = 0; (B, D) show the corresponding plots for *β* = 1. The histograms use 10^4^ independent endpoint samples at steady state. Parameter choices are the same as in Fig. 3B.

### D. Inactive protein controls the switching rates

Although inactive proteins may not play any direct role in gene regulation, they can influence switching between active and inactive states of proteins. Here, we consider that one of the switching rates is a nonlinear function of the inactive-protein level, whereas the other rate remains independent of protein levels. When the inactive protein controls sequestration with a rate

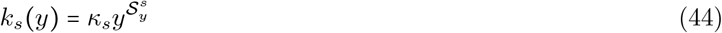

and the dissociation rate *k*_*d*_ is constant, LNA yields a closed-form Fano factor in the fast-switching limit, provided that the system is monostable. The conditions for monostability are obtained from the corresponding deterministic model in Appendix C 2, where the steady states are denoted by 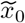 and 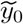 for the active and inactive proteins, respectively. Here, 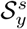 is the logarithmic sensitivity of the sequestration rate *k*_*s*_(*y*) to the level of inactive protein and is defined at steady state as:

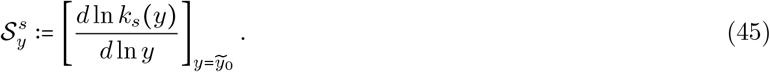

For *β* = 0, a unique and locally asymptotically stable positive steady state exists when 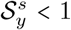. The active- and inactive-protein concentrations have the following closed forms:

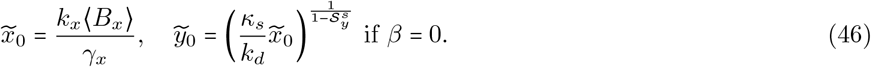

Here, the steady-state active-protein concentration is identical to that in the absence of sequestration.

The *β* = 1 case does not, in general, admit any closed-form expressions for the steady states. Nevertheless, monostability is ensured for 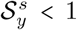, independent of other parameters. The steady-state inactive-protein concentration obeys

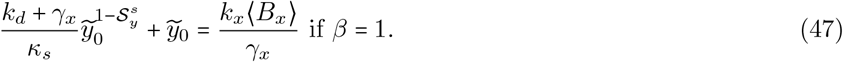

For 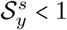, the left-hand side of Eq. (47) is continuous on [0, ∞) and strictly increasing on (0, ∞ ). Its limiting values are 0 as 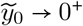 and ∞ as 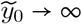. Since *k*_*x*_⟨*B*_*x*_⟩/*γ*_*x*_ > 0, Eq. (47) admits a unique positive root 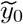. The corresponding active-protein concentration is obtained from

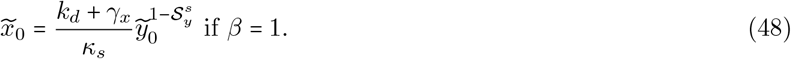

Thus, for 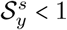, the system admits a unique positive steady state which is locally asymptotically stable. The stability analysis is presented in Appendix C 2.

Upon expansion around the deterministic stable steady states 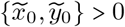, the linearized sequestration propensity becomes

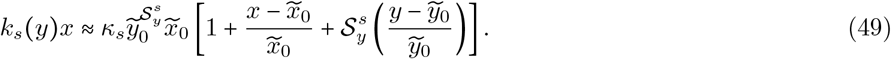

We use this linearized sequestration propensity in the moment dynamics equation [88–90] to calculate the required moments at steady state. The steady-state mean protein levels are

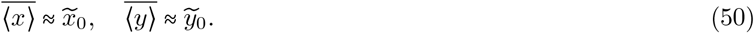

For geometrically distributed burst sizes 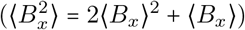, the Fano factor expressions in the fast-switching limit admit the following forms:

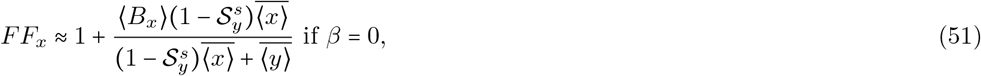

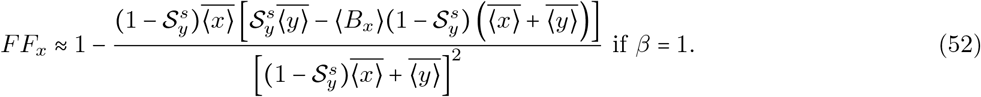

Next, we turn to the opposite situation, where the reversible active-to-inactive switching involves a constant sequestration rate *k*_*s*_ and inactive-protein-dependent dissociation rate

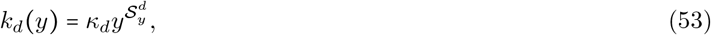

where 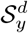 is the logarithmic sensitivity of the dissociation rate to the inactive-protein level. The conditions for a monostable system are discussed in Appendix C 3, where the corresponding deterministic model is presented. 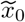 and 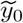 denote the steady-state concentrations of the active and inactive proteins, respectively. The steady-state sensitivity is defined as follows:

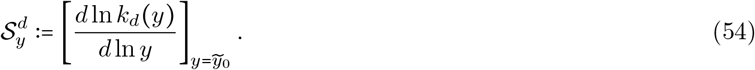

For *β* = 0, a unique positive and locally asymptotically stable steady state exists when 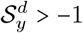. The steady-state protein concentrations have the following forms:

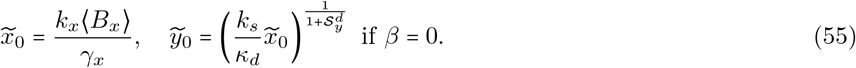

Similar to the previous case, 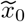 is identical to the steady-state active-protein concentration in the absence of sequestration.

For the *β =* 1 case, 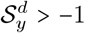 ensures the existence of a monostable system independent of other parameters. The steady-state inactive-protein concentration satisfies

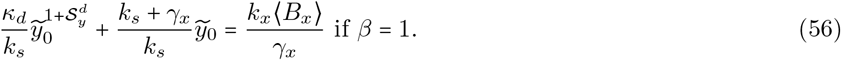

For 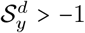 or equivalently 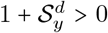, the left-hand side of Eq. (56) is continuous on [0, ∞), strictly increasing on (0, ∞), approaches 0 as 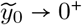, and diverges as 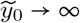. Since *k*_*x*_⟨*B*_*x*_⟩/*γ*_*x*_ > 0, there is exactly one positive solution 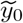. Consequently, 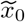 is also uniquely determined, and turns out to be positive:

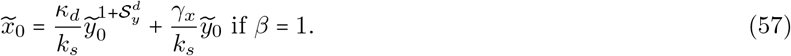

The steady state is locally asymptotically stable for 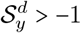. The stability analysis is presented in Appendix C 3. Next, we linearize the nonlinear dissociation propensity around the steady-state inactive-protein level 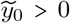 as follows:

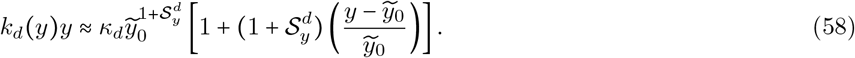

We use this linearized dissociation propensity in the moment dynamics equation [88–90] to evaluate the required steady-state moments. The steady-state mean protein levels are

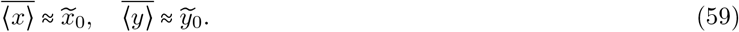

The closed-form Fano factors in the fast-switching limit for geometrically distributed burst sizes 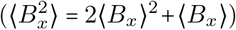 are as follows:

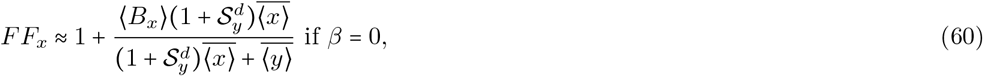

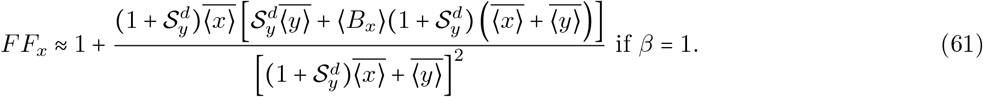

Eqs. (60) and (61) show increasing trends in the active-protein fluctuations with an increasing sensitivity of the dissociation rate to the inactive-protein level 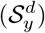. Exactly the opposite trends are seen in Eqs. (51) and (52). The symmetry in the Fano factor with respect to sensitivity can be verified by replacing 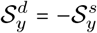 in Eqs. (60) and (61), which then produce Eqs. (51) and (52), respectively.

Connecting back to the results in Sec. III A, the following Fano factor expressions (in the fast-switching limit) shed light on the active-protein fluctuations when the protein is synthesized in non-bursty reactions (*B*_*x*_ = 1 with_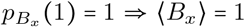_ and 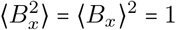. When the sequestration rate depends on the inactive-protein level and the dissociation rate is constant, we have

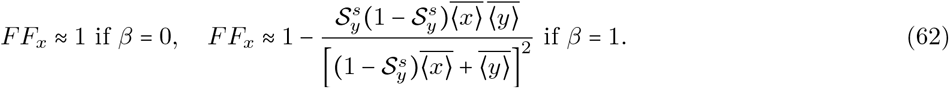

Similarly, with an inactive-protein-dependent dissociation rate and a constant sequestration rate, the Fano factor is reduced to

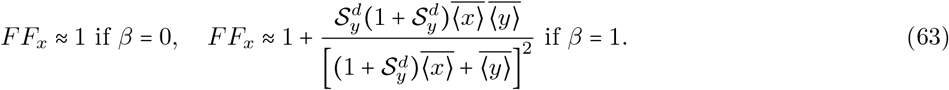

In the above expressions, *FF*_*x*_ ≈ 1 reaffirms our findings in Sec. III A for an inactive protein protected from decay (*β* = 0): the Poisson nature of the active-protein distribution from the non-bursty gene expression remains unaffected by the sequestration kinetics when the switching rates depend only on the inactive-protein level. Thus, for non-bursty synthesis and *β =* 0 to produce Poissonian fluctuations, it is sufficient that one of the switching rates depends on the inactive-protein level while the other remains constant. On the contrary, this is not the case in general when the inactive protein decays at the same rate as the active protein (*β* = 1). In the fast-switching limit, the special case leading to Poissonian fluctuations from the second expression in Eq. (62) is when 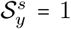, provided the parameter condition for a unique positive stable steady state in Eq. (C22) is satisfied. Similarly, 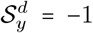, i.e., when the dissociation rate *k*_*d*_ (*y*) ∝ *y* ^−1^, results in Poissonian fluctuations in the fast-switching limit, as evident from the second expression in Eq. (63). In this case, the unique positive stable steady state exists subject to the parameter condition in Eq. (C36).

We normalize the Fano factors from Eqs. (51), (52), (60), and (61) by the Fano factor without sequestration(*FF*_*x*_ = 1 + ⟨*B*_*x*_ ⟩, from Eq. (28)), and plot the Fano factor ratios in Figs. 5A and 5B. In both cases, attenuation of fluctuations is evident as 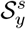 increases and 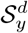 decreases, respectively. Next, we discuss how this framework can be contextualized in two interesting biochemical phenomena.

**FIG. 5.**
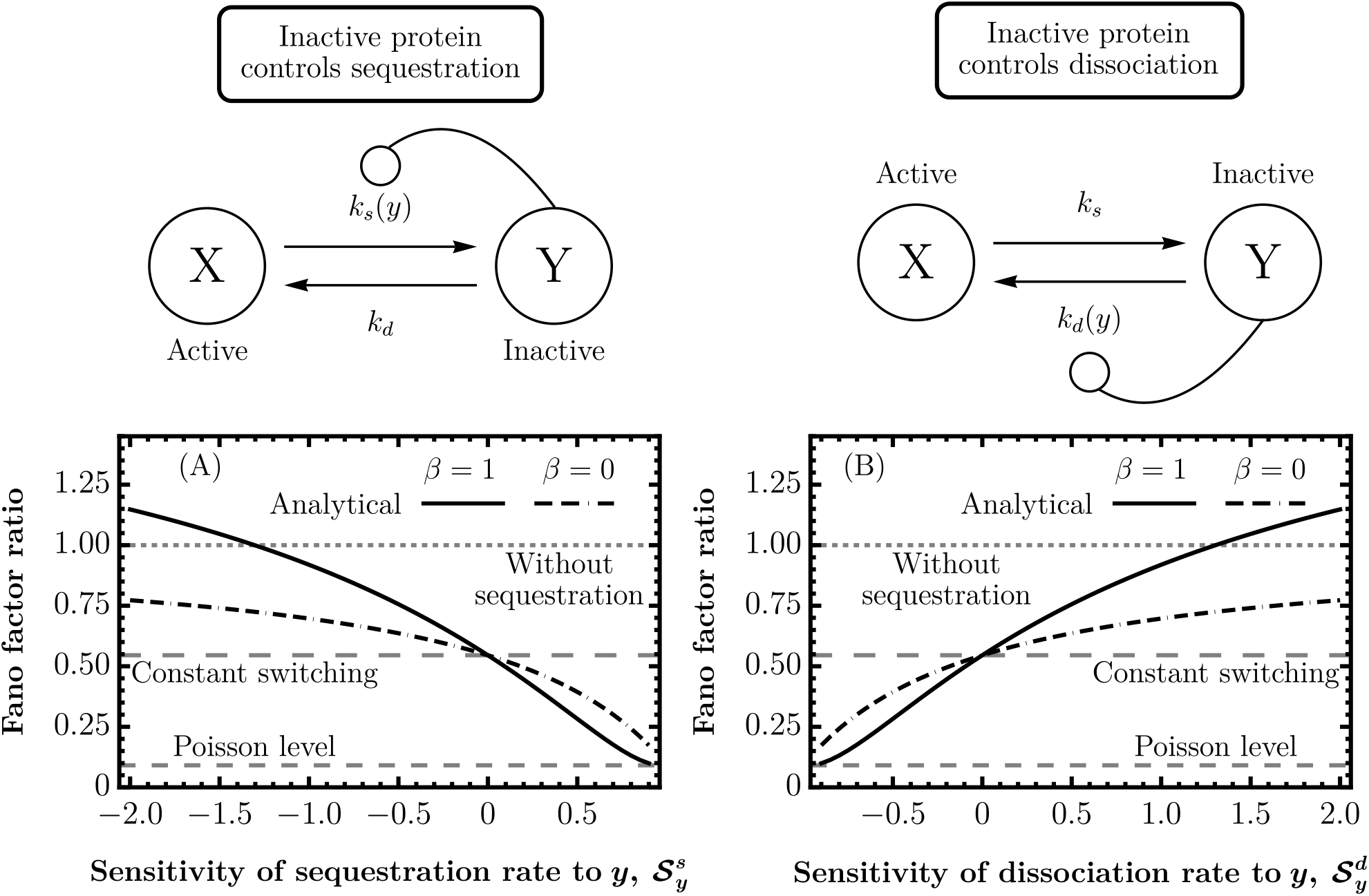
Active-protein fluctuations are attenuated with increasing sequestration-rate sensitivity and decreasing dissociation-rate sensitivity to the inactive-protein level (fast-switching limit). (A) The ratios of the active-protein Fano factors under sequestration, *FF*_*x*_ from Eqs. (51) and (52), to the Fano factor without sequestration, *FF*_*x*_ from Eq. (28), are plotted as functions of 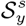. (B) Similarly, the ratios of the active-protein Fano factors under sequestration, *FF*_*x*_ from Eqs. (60) and (61), to the Fano factor without sequestration, *FF*_*x*_ from Eq. (28), are plotted as functions of 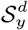. Gray dashed lines indicate the “Without sequestration”, “Constant switching” (the ratio of *FF*_*x*_ in Eq. (27) to *FF*_*x*_ in Eq. (28)), and “Poisson level” (the ratio of *FF*_*x*_ in Eq. (20) to *FF*_*x*_ in Eq. (28)) reference values. The “Constant switching” reference corresponds to the “Fast-switching limit” line in Fig. 2. Parameters: 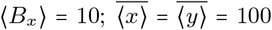 copies; *γ*_*x*_ = 1 and *γ*_*y*_ = 0 for *β* = 0; and *γ*_*x*_ = *γ*_*y*_ = 1 for *β* = 1. Other parameters are adjusted to keep the steady-state protein levels fixed as the sensitivities are varied.

The inactive-protein-dependent sequestration considered in Fig. 5A can be realized through the binding of active proteins to genomic decoy sites. The sequestration rate has the form

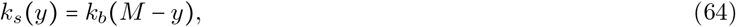

whereas the dissociation rate *k*_*d*_ is constant. Here, *M* is the total number of genomic decoy sites, with *y* ∈ {0, 1, …, *M*} denoting the number of bound proteins. The rate parameter *k*_*b*_ is the propensity coefficient for binding an active protein to one of the *M* − *y* free decoy sites. Hence, the sequestration propensity becomes *k*_*b*_ (*M* − *y*)*x* [62, 104]. At the fully occupied boundary *y = M*, the sequestration rate vanishes, thereby preventing transitions to states with more than *M* occupied sites.

The logarithmic sensitivity is evaluated at an interior steady state, 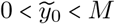, where 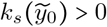. Using 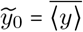 in Eq. (45), the steady-state sensitivity of the sequestration rate to the inactive-protein level is

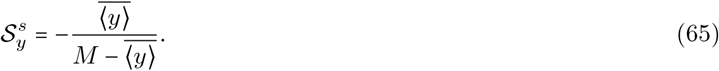

Thus, the sensitivity is negative at every interior operating point. For fixed inactive-protein level 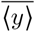, increasing the number of free decoy sites, 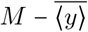, makes the sensitivity less negative and drives it towards zero. As shown in the negative-sensitivity domain of Fig. 5A, sequestration through genomic decoy binding can maintain active-protein fluctuations below the “Without sequestration” level for both values of the bound-protein decay rate considered here, when the bound proteins are protected from decay (*β* = 0) than when they decay at the same rate as the active proteins except at sufficiently negative sensitivities for *β =* 1. However, the Fano factor ratios show that attenuation is stronger (*β* = 1). Protection from decay allows the bound-protein pool to act as a stabilized reservoir. When bound proteins also decay, the additional stochastic loss events and the synthesis needed to maintain the prescribed steady-state protein levels increase fluctuations in the free-protein pool, consistent with previous findings [62].

As a second example, we consider a simplified phase-separation scenario in which the total volume occupied by the inactive protein molecules is much smaller than the system volume and the active-protein level is high. Under these limiting conditions (see Eq. (6)), the sequestration rate becomes a constant (*k*_*s*_), and the dissociation rate takes the form

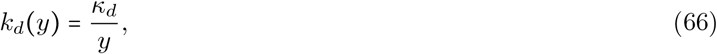

where *κ*_*d*_ is a constant parameter. For *y* > 0, the dissociation reaction occurs with a constant nonzero propensity *k*_*d*_(*y*)*y* = *κ*_*d*_, whereas at *y* = 0 the dissociation propensity is defined to be zero because no inactive protein is available for dissociation. From Eq. (54), the sensitivity is 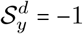.

When the probability of the boundary state *y* = 0 is negligible, the dissociation propensity is effectively constant, and a closed-form approximation for the Fano factor can be obtained when the inactive protein decays at the same rate as the active protein (*β* = 1). As noted in Eq. (C36), the corresponding deterministic system is monostable subject to the constraint *k*_*s*_ *k*_*x*_ ⟨*B*_*x*_⟩ *> κ*_*d*_ *γ*_*x*_ . In this boundary-negligible regime, the steady-state mean protein levels are well approximated by the corresponding deterministic concentrations, 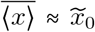 and 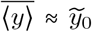. Using standard moment-dynamics methods [88–90], the steady-state moment equations yield:

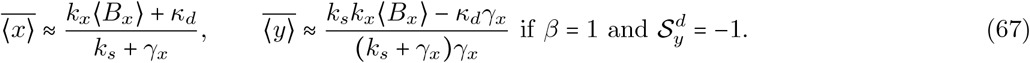

For a geometric burst-size distribution 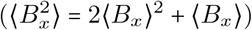, the corresponding Fano factor is given by

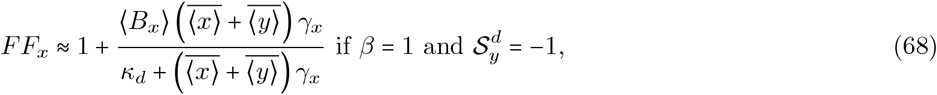

which approaches the Poisson level (*FF*_*x*_ ≈ 1) in the fast-switching limit.

In Fig. 6A, we show the Fano factor ratio in the fast-switching limit for *β =* 1 using Eqs. (61) and (28) when the proteins are synthesized in bursts. The non-bursty case is presented in Fig. 6B using the second expression in Eq. (63). Here, as expected, in the absence of sequestration, the Fano factor is at the Poisson level, *FF*_*x*_ = 1, as shown in Eq. (20). For bursty protein synthesis, decreasing 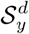 from positive values generally attenuates the fluctuations. In the non-bursty case, however, the dependence is nonmonotonic: the Fano factor ratio reaches a local minimum between 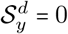 and 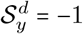 before returning to the Poisson level predicted by the LNA at 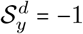. This attenuation is further illustrated by the active-protein time courses and histograms in Fig. 7. This fluctuation-attenuating feature, emerging from our general sequestration model, is in qualitative agreement with previous studies of phase separation and noise buffering [67, 69].

**FIG. 6.**
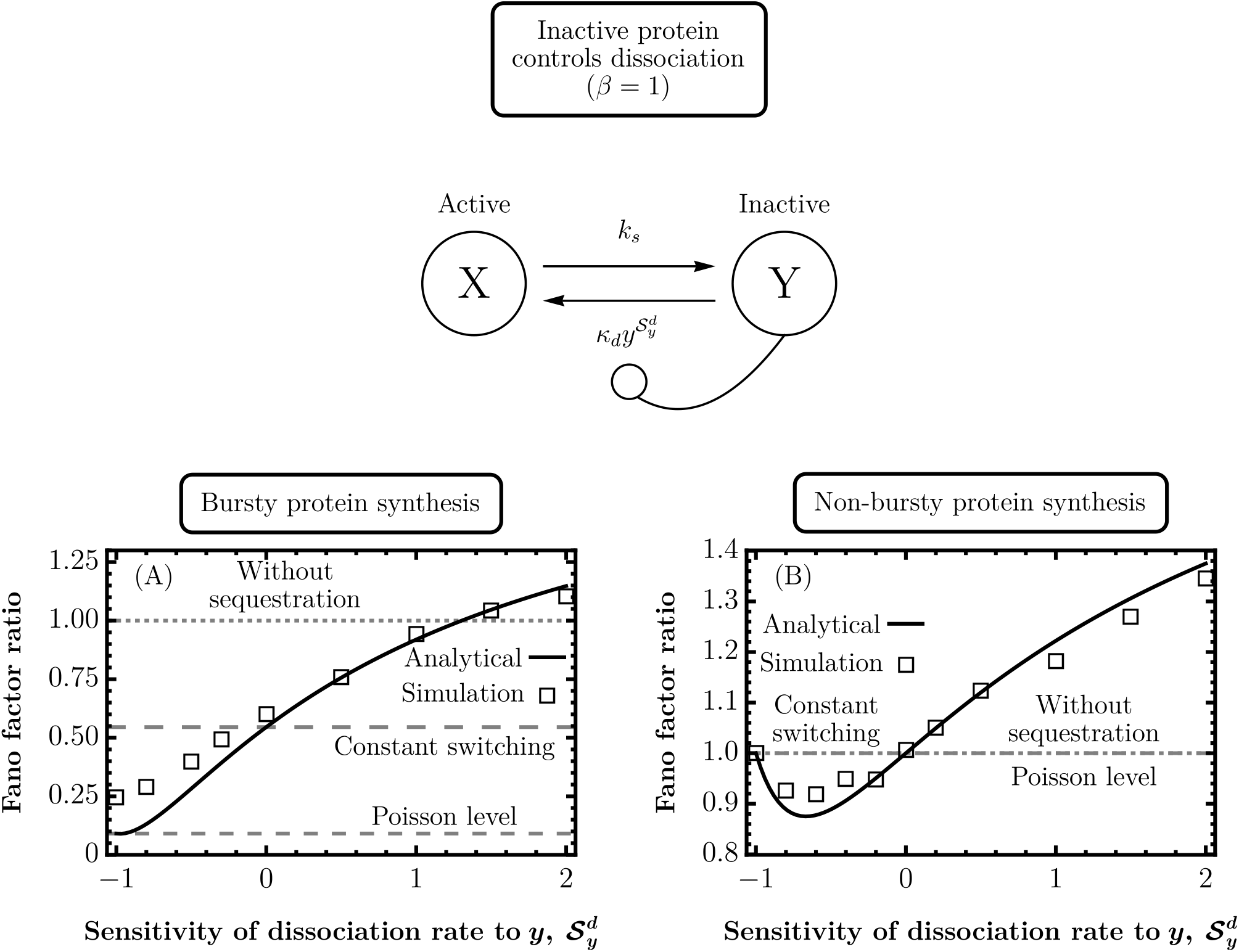
Active-protein fluctuations are attenuated with decreasing dissociation-rate sensitivity to the inactive-protein level for bursty synthesis but vary nonmonotonically for non-bursty synthesis when both protein species decay at the same rate (*β =* 1). (A) The ratio of the active-protein Fano factor under sequestration, *FF*_*x*_ from Eq. (61), to the Fano factor without sequestration, *FF*_*x*_ from Eq. (28), is plotted as a line for bursty synthesis. (B) The corresponding non-bursty ratio is plotted as a line using the second expression in Eq. (63) and the Fano factor without sequestration, *FF*_*x*_ from Eq. (20). Symbols show stochastic-simulation estimates [98, 99] from 10^4^ independent endpoint samples at steady state. Here, 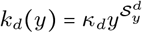 *k*_*s*_ is constant. In (A), dashed lines mark the “Without sequestration”, “Constant switching” (the “Fast-switching limit” line in Fig. 2), and “Poisson level” (the ratio of *FF*_*x*_ in Eq. (20) to *FF*_*x*_ in Eq. (28)) reference values; all three ratios coincide in (B). Parameters: ⟨*B*_*x*⟩_ *=* 10 in (A); 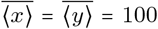 copies; and *γ*_*x*_ *= γ*_*y*_ *=* 1. Analytical lines represent the fast-switching limit; stochastic simulations use the finite switching rate 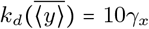. Other rates are adjusted to maintain the steady-state protein levels.

**FIG. 7.**
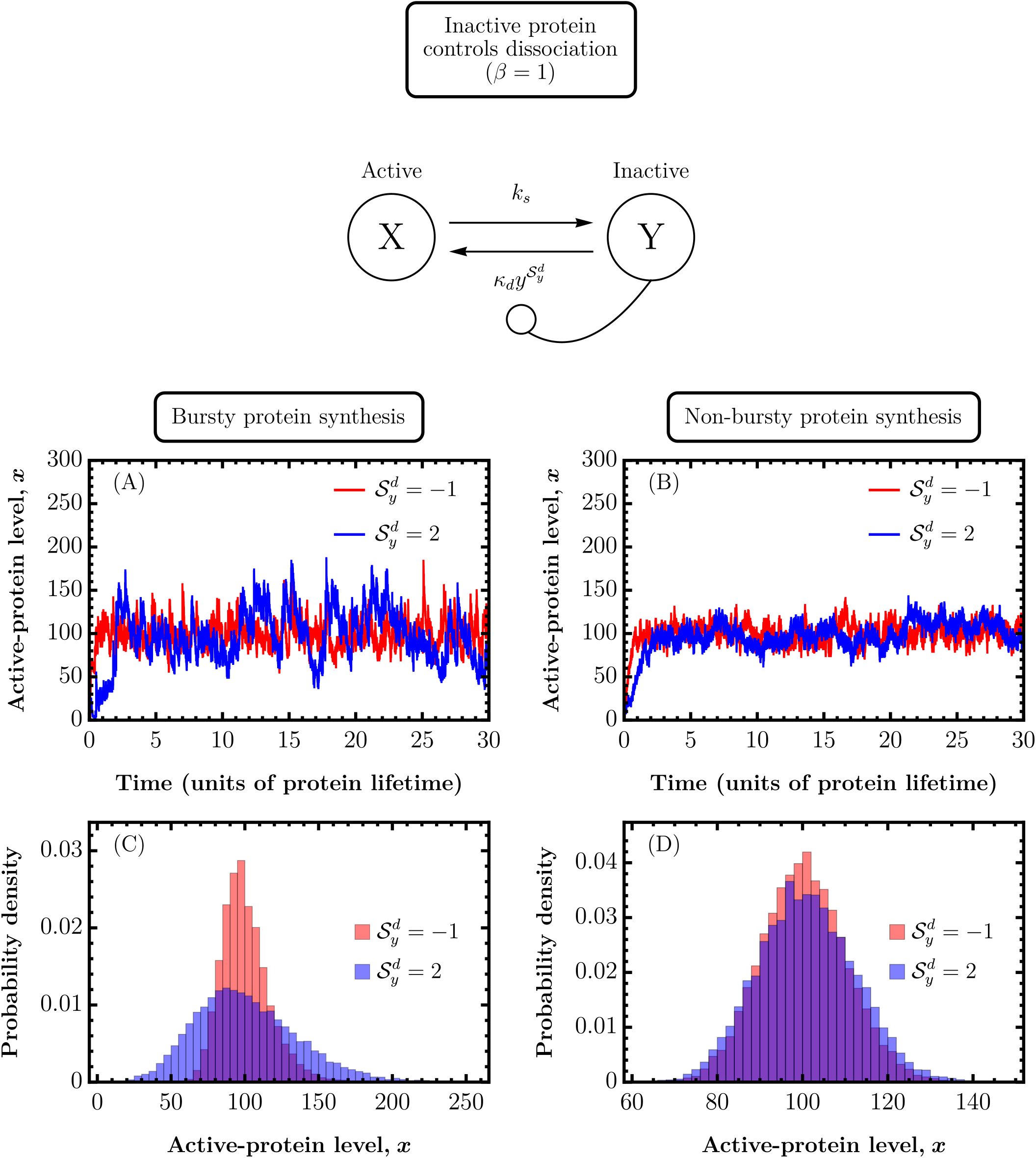
Active-protein fluctuations are attenuated at lower dissociation-rate sensitivity to the inactive-protein level when both protein species decay at the same rate (*β =* 1; continued from Fig. 6). The dissociation rate is 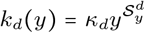, while the sequestration rate *k*_*s*_ is constant. (A, C) Sample trajectories and probability density histograms, respectively, from stochastic simulations [98, 99] for different values of 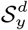 under bursty synthesis; (B, D) show the corresponding non-bursty results. The histograms use 10^4^ independent endpoint samples at steady state. Parameter choices are the same as in Fig. 6.

## IV. CONCLUSION

In this article, we investigate the effects of different types of reversible sequestration mechanisms on the intracellular variability of proteins synthesized in bursts and decaying following a first-order process. Our stochastic framework models the sequestration and dissociation of a single protein molecule as active-to-inactive and inactive-to-active switching reactions, respectively. The switching rates can be arbitrary functions of the free (active) or sequestered (inactive) protein levels. Furthermore, we consider two scenarios for the decay of inactive protein: when it is protected from decay and when it decays at the same rate as the active protein.

For the special case where the protein synthesis is non-bursty, the switching rates depend only on the inactive-protein level, and the inactive protein is protected from decay, our exact result shows that the active-protein distribution is Poisson. In other words, fluctuations in the active-protein level remain unaffected by the sequestration kinetics. This invariance in fluctuation profile holds even when more than one protein molecule cooperatively undergoes sequestration. This result is significant for the inactivation of biomolecules through complex formation, e.g., in ribosomal downregulation in bacteria [73, 74], nonspecific binding of transcription factor proteins to genomic “decoy” sites [62, 63], and for low-abundance proteins, for which the total noise is dominated by the intrinsic noise [11, 105].

For sequestration kinetics with constant switching rates, we compare the Fano factor with respect to the Fano factor in the absence of sequestration by taking their ratios (see Fig. 2). The steady-state protein levels are kept fixed. To simplify the general results, we consider protein burst sizes that are geometrically distributed. For both protected and decaying inactive protein, the active-protein fluctuations under sequestration remain below the fluctuation level in the absence of sequestration for positive dissociation rates. Notably, when the inactive protein decays at the same rate as the active protein, fluctuations are enhanced compared to the case when the inactive protein is protected from decay. However, an increasing dissociation rate attenuates active-protein fluctuations in both cases.

To probe general mechanisms with the sequestration rate nonlinearly dependent on the active-protein level and a constant dissociation rate, we use the LNA and obtain closed-form Fano factor expressions in the fast-switching limit, considering geometrically distributed burst sizes. For fixed steady-state protein levels, we plot the Fano factor ratio with the logarithmic sensitivity of the sequestration rate. We restrict the sensitivity to a biologically plausible range spanning both positive and negative values that ensures a monostable regime. We observe that an increasing sensitivity attenuates active-protein fluctuations for both protected and decaying inactive protein (see Fig. 3A). With the support of stochastic simulations, we demonstrate this attenuation in the post-translational modification of proteins, which follows the Michaelis-Menten kinetics. Fluctuations are attenuated as the enzyme becomes less saturated and the maximal inactivation capacity of the enzyme increases (see Fig. 3B and Fig. 4). The active protein pool fluctuates more when the inactive protein decays at the same rate as the active protein than when it is protected from decay. Interestingly, for bursty protein synthesis, enzymatic inactivation cannot attenuate active-protein fluctuations below the level corresponding to constant switching rates in the fast-switching limit. By contrast, for non-bursty protein synthesis, this attenuation is limited at the Poisson level [100].

Similar effects are seen where the switching rates become arbitrary nonlinear functions of the inactive-protein level. For an inactive-protein-dependent sequestration rate and a constant dissociation rate, active-protein fluctuations are attenuated with increasing sensitivity (see Fig. 5A). An example of this is found in the context of genomic decoy binding, where a growing number of free decoy sites raises the sensitivity of the sequestration rate and dampens fluctuations. Previous studies showed that decoy binding reduces gene expression noise in autoregulatory motifs [62, 63, 78]. This reduction is pronounced when the decoy-bound proteins are protected from decay [62]. These features emerge naturally from our results shown in Fig. 5A (see the negative sensitivity domain), which are based on a generalized inactive-protein-dependent sequestration rate and do not consider specific genetic architectures specialized in regulating noise. Interestingly, both enzymatic inactivation of proteins (see Fig. 3B and Fig. 4) and inactivation via decoy binding (see the negative sensitivity domain in Fig. 5A) show more fluctuations when inactive proteins are unprotected (*β =* 1) than when they are protected (*β =* 0) from decay. This is also the case in sequestration with constant switching rates (see Fig. 2). This similarity plausibly points towards a more general connection between active-protein fluctuations and the decay of inactivated proteins.

The effect of increasing sensitivity on the active-protein fluctuations for an inactive-protein-dependent dissociation rate and a constant sequestration rate (see Fig. 5B) is just the opposite of what is found in Fig. 5A. Interestingly, the LNA shows that for non-bursty protein synthesis and protected inactive protein, fluctuations are Poissonian whenever any one of the switching rates exclusively depends on the inactive-protein level. When the inactive protein shares the same decay rate as the active protein, the LNA predicts approximately Poissonian fluctuations in the fast-switching limit if the dissociation rate varies inversely with the inactive-protein level. This switching mechanism is similar to intracellular phase separation, in the limiting case where the protein level in the dilute phase is high, and the total volume of protein molecules in the droplet phase is much smaller than the system volume. Our analysis stemming from a general stochastic reaction scheme is in qualitative agreement with the noise buffering observed during phase separation [67]. However, this also emphasizes that the noise attenuation can be observed for a broad class of biochemical processes adhering to a similar type of switching condition and therefore cannot be construed as an exclusive feature of biomolecular condensates. Interestingly, a recent theoretical study based on nonequilibrium considerations has successfully disentangled noise attenuation from concentration buffering, which is often linked to phase-separating systems [69].

Our present analysis is restricted to analyzing the effects of regulated sequestration and dissociation rates on active-protein fluctuations. The analytical results presented here are based on the LNA under the assumption of small noise around a unique stable steady state [95]. For strongly nonlinear switching propensities, low-copy-number regimes, and parameter choices producing multistability or multimodal protein distributions, the conventional LNA may become inaccurate or fail to capture the system behavior [106]. Similarly, the use of non-elementary propensities, such as coarse-grained Hill-type forms, in stochastic simulations should be regarded as an approximation whose validity depends on the underlying biochemical mechanisms and the separation of timescales [107–109]. The stochastic framework can be extended to include an autoregulated protein-synthesis rate (assumed constant here) and feedback from the inactive protein [110]. One can also relax the constraint of fixed steady-state protein levels considered here and explore different biologically plausible regimes of active- and inactive-protein levels while keeping the total protein level fixed. Recent papers have gone beyond expression noise and explored variability in cellular timekeeping [111–115]. The present framework can be extended in this direction to investigate the role of reversible sequestration in maintaining precision in biochemical clocks [116]. Another interesting extension is to incorporate extracellular or global fluctuations directly in our model by considering the protein synthesis rate as a deterministic [117] or stochastic [67] representation of the environmental signal. This will introduce extrinsic fluctuations into the system, in addition to those inherent in the stochastic synthesis and degradation of proteins. It will be a nontrivial problem to discern the biochemical mechanisms that can attenuate different types of fluctuations [118]. Furthermore, the question of fluctuation propagation to downstream processes [76] from the active protein may reveal hitherto unknown architectural features of regulatory cascades [57, 119, 120], which are ubiquitous in both sensory and developmental decision-making networks in eukaryotes [121].

## ACKNOWLEDGMENTS

The authors thank Christoph Zechner for helpful discussion. AS acknowledges the support of NIH-NIGMS via grant R35GM148351.

## Appendix A Sequestration with non-bursty synthesis, protein multimerization, and protected inactive complexes

Here, we consider a special case in which protein synthesis is non-bursty, with *B*_*x*_ *=* 1 and 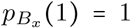, and sequestration occurs through cooperative multimerization. In each sequestration event, *n* active protein molecules form one inactive complex. The inactive complexes are protected from decay (*γ*_*y*_ = 0), and their abundance controls the sequestration and dissociation rates. Accordingly, *x* (*t*) denotes the number of active protein molecules, whereas *y*(*t*) denotes the number of inactive complexes. The corresponding chemical kinetics are obtained by modifying those in Table I. The sequestration propensity is obtained by counting the distinct combinations of *n* active protein molecules, among a total of *x*, that can cooperatively form an inactive complex:

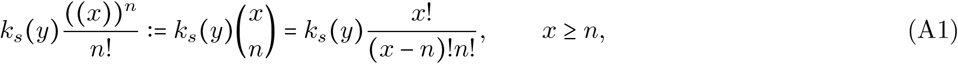

where

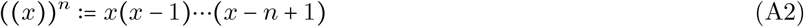

is the falling factorial with ((*x*)) ^*n*^ 0 for *x < n*. Hence, for sequestration of monomers (*n =* 1), the propensity is simply *k*_*s*_ (*y*)*x* and for dimers (*n* = 2) it is *k*_*s*_ (*y*)*x*(*x* − 1)/2. The modified chemical kinetics are presented in Table III.

**TABLE III.** Chemical kinetics for non-bursty protein synthesis and cooperative multimerization when the inactive complexes are protected from decay. Proteins are synthesized at rate *k*_*x*_. In each sequestration event, *n* active protein molecules are sequestered cooperatively to form one inactive complex. Here, *x* (*t*) denotes the number of active protein molecules, whereas *y*(*t*) denotes the number of inactive complexes. The inactive complexes are protected from decay (*γ*_*y*_ = 0). The sequestration and dissociation rates, *k*_*s*_(*y*) and *k*_*d*_(*y*), respectively, are arbitrary functions of the inactive-complex count *y*(*t*). For *n* = 1, this model reduces to a special case of the general model in Table I.

| Biochemical reactions | Changes in species counts | Probability of reaction in the time interval $(t, t + dt]$ |
| --- | --- | --- |
| Protein synthesis | $x \mapsto x + 1$ | $k_x dt$ |
| Sequestration (active-to-inactive state switching) | $x \mapsto x - n$<br>$y \mapsto y + 1$ | $k_s(y) \frac{((x))^n}{n!} dt$ |
| Dissociation (inactive-to-active state switching) | $x \mapsto x + n$<br>$y \mapsto y - 1$ | $k_d(y) y dt$ |
| Decay of active protein | $x \mapsto x - 1$ | $\gamma_x x dt$ |

The chemical master equation governing the time evolution of the joint probability distribution *p* (*x, y, t*) of the active-protein molecule count and the inactive-complex count can be written as [72]

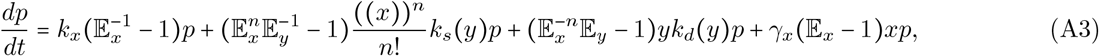

with the following definitions of the van Kampen step operators applied to an arbitrary function *ϕ x, y* [95]:

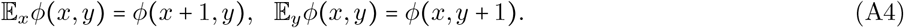

The corresponding inverse step operators naturally become

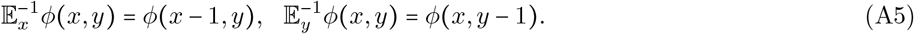

The *n*th-order extensions read as

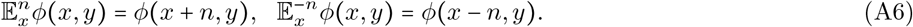

Rearranging the step operators, Eq. (A3) becomes

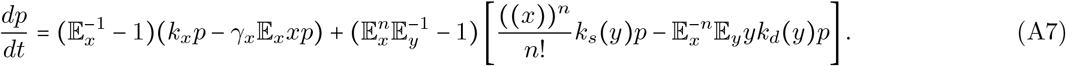

To construct a stationary solution of Eq. (A7), we seek a distribution *p* (*x, y*) = lim_*t*→∞_ *p* (*x, y, t*) satisfying the following sufficient pairwise-balance conditions:

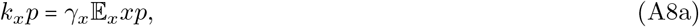

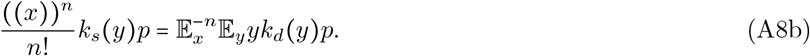

To solve Eqs. (A8a) and (A8b), we seek a separable form for the joint distribution [82–85]:

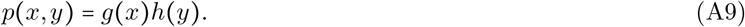

By inserting Eq. (A9) into Eq. (A8a) and dividing by *h y* we obtain a first-order recursion relation for *g* (*x*) :

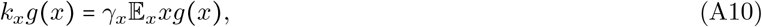

Similarly, inserting Eq. (A9) into Eq. (A8b), we get

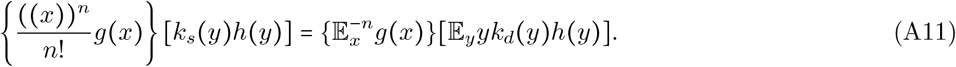

Next, we solve the recursion in Eq. (A10) iteratively to obtain the steady-state distribution *g* (*x*) of the active-protein count. The marginal active-protein distribution at steady state,

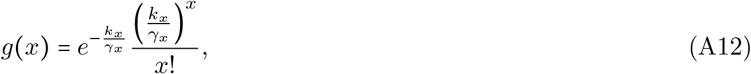

is Poisson with the mean count

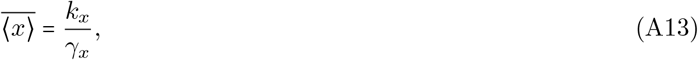

and thus remains unaffected by the sequestration process. Inserting Eq. (A12) into Eq. (A11) and simplifying yields a first-order recursion relation for *h* (*y*) :

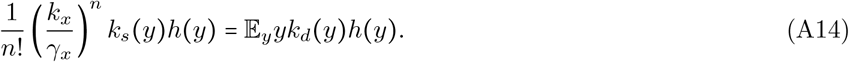

The marginal distribution of inactive multimeric complexes is obtained by solving Eq. (A14) and takes the form:

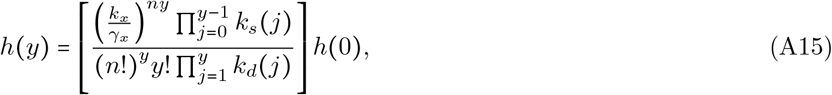

where *h* (0) is determined by the normalization condition:

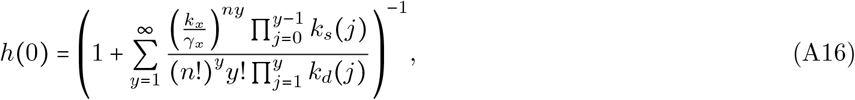

provided that the normalization sum converges. Eq. (A15) is mathematically analogous to the stationary distribution of a one-species birth–death process for the inactive complexes, with effective complex-formation propensity (*k*_*x*_ */ γ*_*x*_)^*n*^*k*_*s*_ (*y*) *n*! and complex-removal propensity *yk*_*d*_ (*y*) . Equivalently, *k*_*d*_ (*y*) is the per-complex removal rate. In the present biochemical interpretation, these effective birth and death events correspond to multimeric sequestration and dissociation, respectively, rather than to synthesis and decay of the complexes. The cooperativity index *n* controls the dependence of the effective complex-formation propensity on the mean active-protein count, 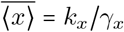. In the special case in which both *k*_*s*_(*y*) = *k*_*s*_ and *k*_*d*_(*y*) = *k*_*d*_ are constant, Eq. (A15) reduces to a Poisson distribution with mean inactive-complex count

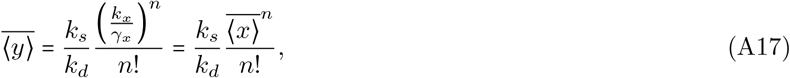

where in the last step, we have used Eq. (A13). Because each inactive complex contains *n* protein molecules, the mean number of protein molecules in the inactive state is 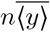.

## Appendix B Statistical moments for constant switching rates

The time evolution of the generalized moment ⟨*x*^*m*^*y*^*l*^⟩ with non-negative integers *m* and *l* can be described using the standard technique of moment dynamics [88–90]. We turn our attention to the reaction kinetics in Table I for the special case when the sequestration and dissociation rates are independent of protein levels. These switching rates are indicated by *k*_*s*_ and *k*_*d*_, respectively. The time evolution of the generalized moment in this case follows:

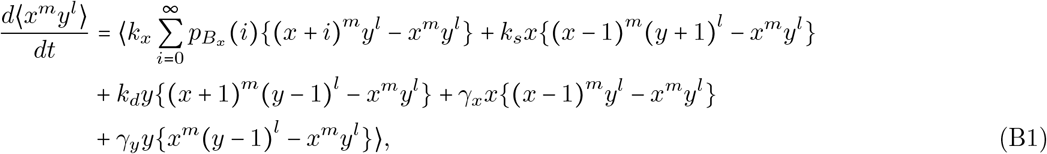

where 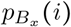 is an arbitrary probability mass function for the burst size *B*_*x*_ as defined in Eq. (1). Upon expansion at steady state, Eq. (B1) produces the following set of coupled algebraic equations involving 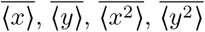, and 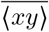:

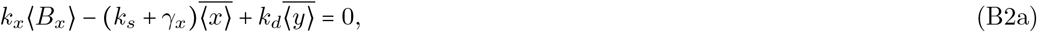

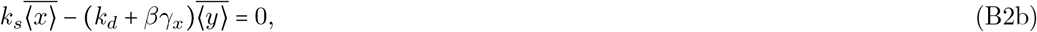

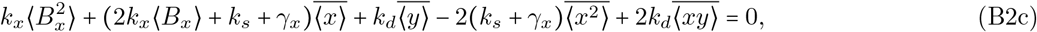

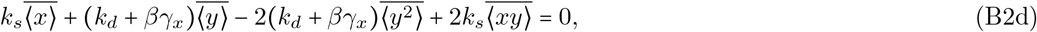

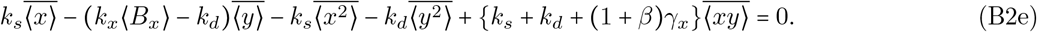

Here, ⟨*B*_*x*_⟩ and 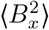 are the first- and second-order moments of the burst-size distribution with the respective definitions:

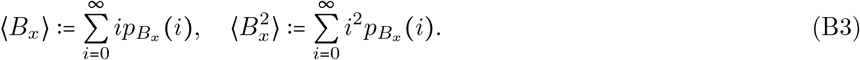

The relative decay rate of the inactive protein *β = γ*_*y*_ */ γ*_*x*_ is zero when it is protected from decay (*γ*_*y*_ *=* 0) and becomes 1 when it has the same decay rate as the active protein (*γ*_*x*_ = *γ*_*y*_ ).

To compute the Fano factor of the active-protein distribution, we need the expressions of 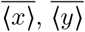, and 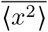 which come from solving Eqs. (B2a)–(B2e):

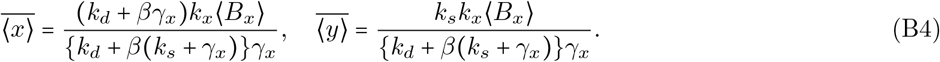

Finally, the second-order moment is obtained as

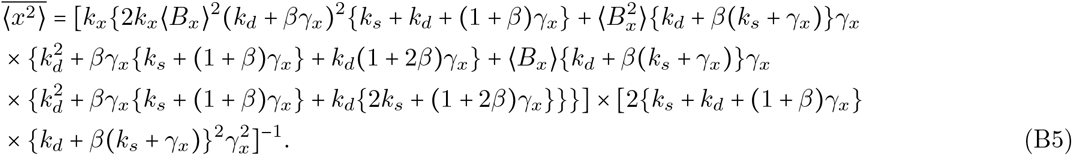

We take the expression of the mean active-protein level from Eq. (B4) and that of the second moment from Eq. (B5) and plug them into Eq. (13). Thus, we obtain the exact closed-form Fano factor in Eq. (17) for a general burst size distribution.

## Appendix C Deterministic analysis: conditions for unique stable steady states

In the deterministic formulation of our stochastic model (see Table I), i.e., when the protein abundances are high enough so that fluctuations are negligible, the following reaction rate equations describe the time evolution of the concentrations of active 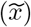 and inactive 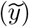 proteins:

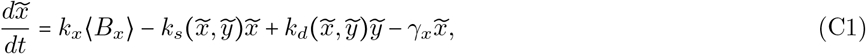

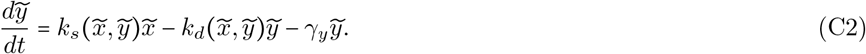

In Eq. (C1), the average burst size 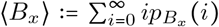, where the burst sizes *B*_*x*_ are distributed according to an arbitrary probability mass function 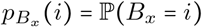 with *i* ∈ {0, 1, 2, …}. Since the effective cellular volume is taken to be unity in our model [96, 122–124], copy numbers and concentrations are numerically identical. Furthermore, the macroscopic rates used in the deterministic description are also numerically identical to their stochastic counterparts. Next, we will consider three different switching cases based on how the sequestration and dissociation rates depend on the active- or inactive-protein concentrations. In each of these cases, we will analyze the conditions for the existence of unique stable steady states for the active and inactive protein species.

### 1. The sequestration rate depends on the active-protein concentration

When the sequestration rate has the form 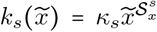 (*κ*_*s*_ is the rate parameter and 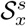 is the logarithmic sensitivity of the sequestration rate to the active-protein concentration) and the dissociation rate *k*_*d*_ is independent of protein concentrations, Eqs. (C1) and (C2) take the following forms at steady state (denoted by subscript 0):

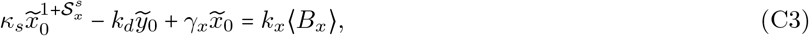

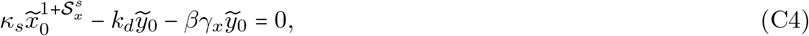

respectively. Here, *β* := *γ*_*y*_/*γ*_*x*_ denotes the relative degradation rate of the inactive protein. Throughout this analysis, we assume *k*_*x*_ ⟨*B*_*x*_⟩ > 0, *γ*_*x*_ > 0, *κ*_*s*_ > 0, and *k*_*d*_ > 0.

When the inactive protein is protected from decay (*β* = 0), it is easy to see from these equations that

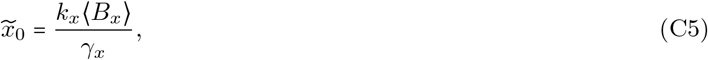

irrespective of the sensitivity 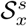. The corresponding steady-state inactive-protein concentration from Eq. (C4) is:

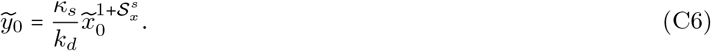

Hence, for *β* = 0 and *k*_*d*_ > 0, a finite and unique positive steady state exists for any 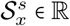.

For *β* = 1, Eqs. (C3) and (C4) show that a unique positive-valued 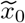 exist whenever 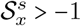, without any further constraints on the other parameters. This can be checked by first extracting 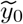 from Eq. (C4) in terms of 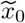 as

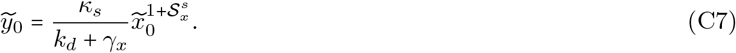

Plugging Eq. (C7) into Eq. (C3) produces

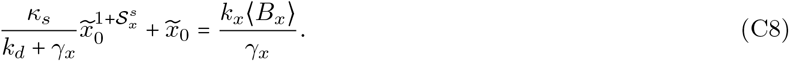

For 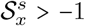, the left-hand side of Eq. (C8) is continuous on [0, ∞) and strictly increasing on (0, ∞). Moreover, it approaches zero as 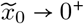 and diverges as 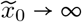. Hence, Eq. (C8) admits a unique positive solution for 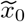. Once this solution is fixed, the uniqueness of 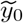 follows from Eq. (C7). At the borderline value 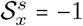, Eq. (C8) gives a unique positive-valued 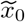 provided (*k*_*d*_ + *γ*_*x*_ )*k*_*x*_ ⟨*B*_*x*_ ⟩ > *κ*_*s*_ *γ*_*x*_ :

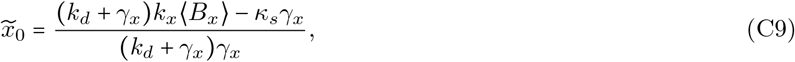

whereas the corresponding unique positive-valued 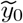 comes from Eq. (C7):

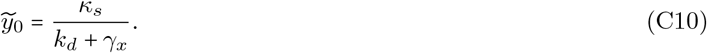

Next, we perform the linear stability analysis [125] on Eqs. (C1) and (C2) with the particular switching rate forms in hand. The conditions for the existence of a locally asymptotically stable steady state are *τ* = *λ*_*x*_ + *λ*_*y*_ < 0 and Δ = *λ*_*x*_*λ*_*y*_ *>* 0, where *λ*_*x*_ and *λ*_*y*_ are the eigenvalues of the Jacobian of the linearized system. *τ* and Δ are the trace and determinant of the Jacobian, respectively. The Jacobian evaluated at the unique steady state reads as follows:

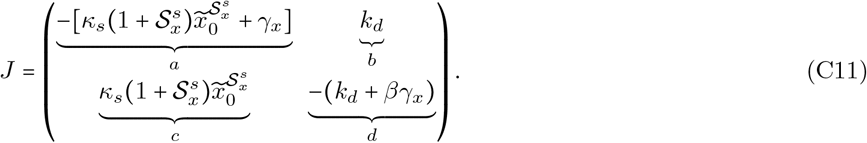

For 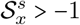, or equivalently 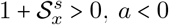 and *c* > 0. Therefore, with *b* > 0 and *d* < 0, the trace becomes

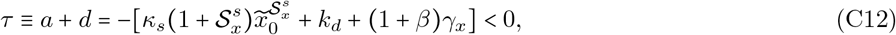

whereas the simplified determinant is

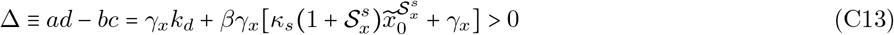

for both *β* = 0 and *β* = 1. For 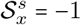, the Jacobian gives:

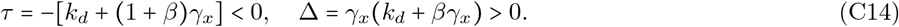

Therefore, the corresponding unique positive steady state is locally asymptotically stable at 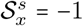 for both *β* = 0 and *β* = 1, whenever it exists.

### 2. The sequestration rate depends on the inactive-protein concentration

Assuming the sequestration rate to be of the form 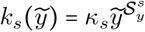 (*κ*_*s*_ is the rate parameter and 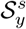 is the logarithmic sensitivity of the sequestration rate to the inactive-protein concentration) and the dissociation rate *k*_*d*_ independent of protein concentrations, we obtain the following equations from Eqs. (C1) and (C2) at steady state:

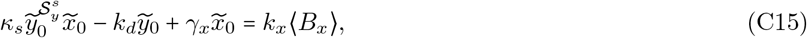

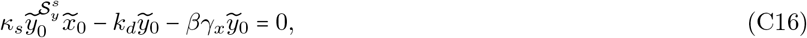

respectively. As mentioned before, *β* := *γ*_*y*_/*γ*_*x*_ denotes the relative degradation rate of the inactive protein. Furthermore, we assume *k*_*x*_ ⟨*B*_*x*_ ⟩ > 0, *γ*_*x*_ > 0, *κ*_*s*_ > 0, and *k*_*d*_ > 0. For the *β* = 0 case, the above equations yield the following unique positive steady state:

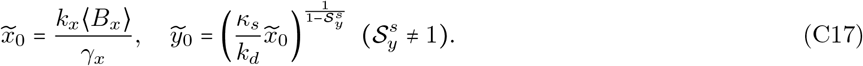

At the borderline value 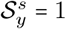, Eq. (C16) reduces to

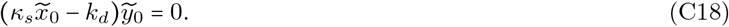

Therefore, either 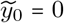 when 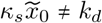 or a continuum of 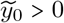 when 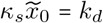. Hence, no unique positive steady state exists at 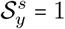 for *β* = 0.

For *β* = 1, it can be shown analytically that the parameter-free condition for unique steady states is 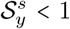. To show the uniqueness, we express 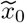 in terms of 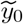 from Eq. (C16):

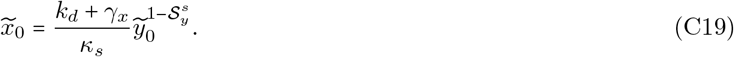

By substituting 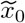 from Eq. (C19) into Eq. (C15), we get

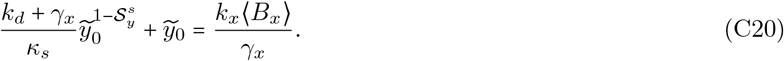

For 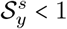 the left-hand side in Eq. (C20) is continuous on [0, ∞), strictly increasing on (0, ∞) with limiting values 0 and ∞ as 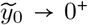 and 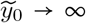, respectively. Therefore, Eq. (C20) admits exactly one positive solution for 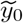. Consequently, a unique positive solution for 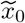 follows from Eq. (C19). The borderline case at 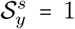 modifies Eq. (C20) into

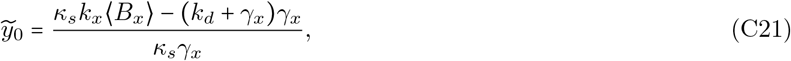

and the condition for unique positive roots turns out to be

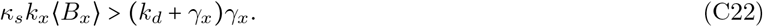

The corresponding unique positive solution for 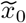 follows from Eq. (C19) with the substitution 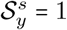:

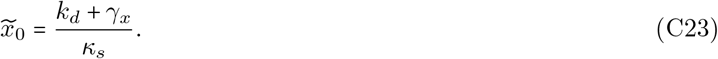

The Jacobian computed from Eqs. (C1) and (C2) with the current sequestration setting and evaluated at the steady state is as follows:

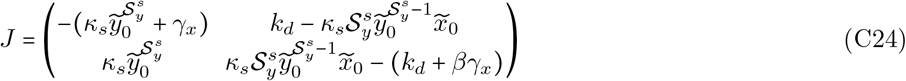

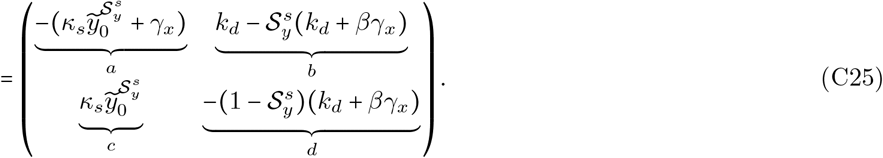

where in the last step we have used 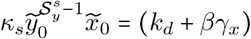 from Eq. (C16). For 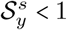, the term *d* < 0, which makes the trace

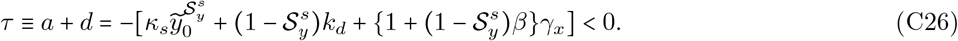

After simplification, the determinant Δ ≡ *ad* − *bc* becomes

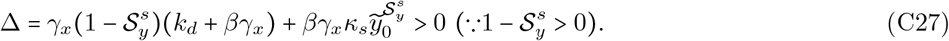

For 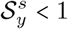, a negative trace and a positive determinant imply that the unique steady state is locally asymptotically stable for both *β* = 0 and *β* = 1. For *β* = 1 and the borderline case at 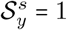, the Jacobian gives

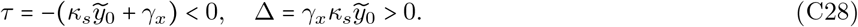

Hence, the steady state is also locally asymptotically stable whenever it exists at 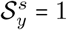 for *β* = 1.

### 3. The dissociation rate depends on the inactive-protein concentration

Here, we assume the dissociation rate to be of the form 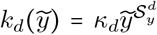 (*κ*_*d*_ is the rate parameter and 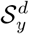 is the logarithmic sensitivity of the dissociation rate to the inactive-protein concentration) and the sequestration rate *k*_*s*_ independent of protein concentrations. At steady state, Eqs. (C1) and (C2) are modified accordingly into

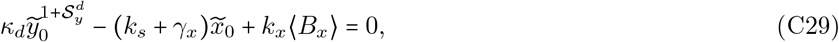

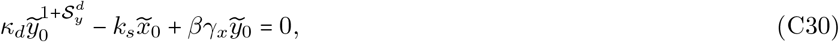

respectively, where *β* := *γ*_*y*_/*γ*_*x*_ is the relative degradation rate of the inactive protein. In our analysis, we assume *k*_*x*_⟨*B*_*x*_⟩ > 0, *γ*_*x*_ > 0, *k*_*s*_ *>*0, *k*_*d*_ *>*0

For *β* = 0, we obtain the following unique positive steady state:

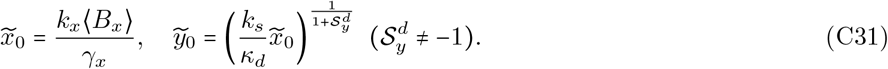

At the borderline case 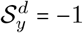. Eq. (C30) reduces to

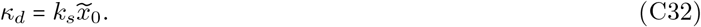

If 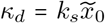. Eq. (C30) admits a continuum of positive-valued 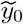, otherwise, no positive steady state exists. Thus, a unique positive steady state does not exist at 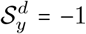 for *β* = 0. If 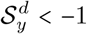, or equivalently 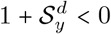, a unique positive steady state exists.

For *β =* 1, a unique steady state is guaranteed when 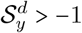, without any further restriction on the other parameters. To verify, we write 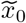 in terms of 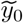 from Eq. (C30) as:

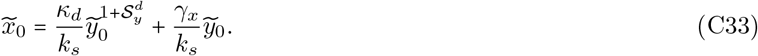

Upon substituting 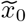 from Eq. (C33) into Eq. (C29), we get

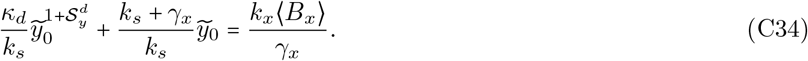

For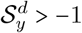, the left-hand side of Eq. (C34) is continuous on [0,∞), strictly increasing on (0, ∞) with limiting values 0 and ∞ as 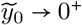 and 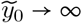, respectively. Since *k*_*x*_ ⟨*B*_*x*_⟩ / *γ*_*x*_ > 0, Eq. (C34) has a unique positive root 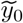. Once 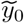 is fixed, a unique positive 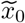 follows from Eq. (C33). For the borderline case, 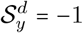, a unique positive 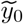 emerges from Eq. (C34):

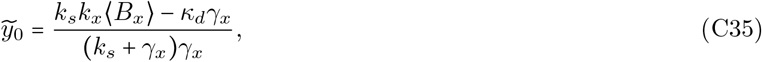

with the condition

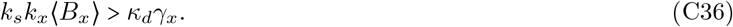

The corresponding unique positive 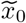 from Eq. (C33) is

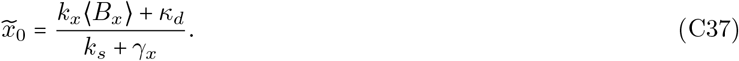

The Jacobian computed from Eqs. (C1) and (C2) with the current dissociation setting and evaluated at the steady state takes the form

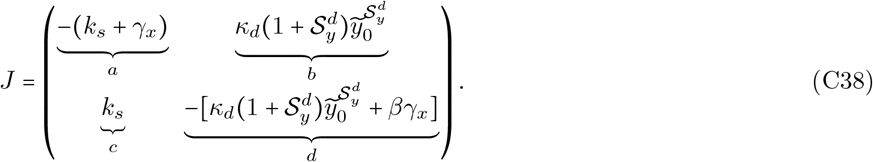

With 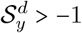 or equivalently 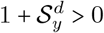, *b* > 0 and *d* < 0 for both *β* = 0 and *β* = 1. Hence, the trace

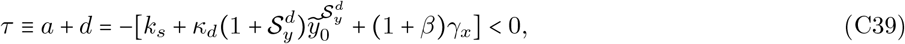

for both *β* = 0 and *β* = 1. Similarly, the determinant in its simplified form

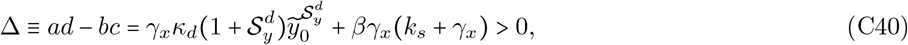

for both *β =* 0 and *β =* 1. For *β =* 0 and 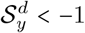, the trace is not sign-definite. However, the determinant becomes negative, implying that the steady state is unstable. Therefore, the unique steady state is locally asymptotically stable when 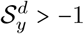 for both *β* values. For the steady state at the borderline case 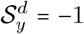 and *β =* 1, the Jacobian gives:

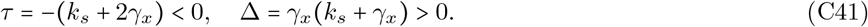

Therefore, whenever this positive steady state exists, it is also locally asymptotically stable.

